# Transcriptional inhibition after irradiation occurs preferentially at highly expressed genes in a manner dependent on cell cycle progression

**DOI:** 10.1101/2023.11.20.567799

**Authors:** Zulong Chen, Xin Wang, Xinlei Gao, Nina Arslanovic, Kaifu Chen, Jessica Tyler

## Abstract

In response to DNA double strand damage, ongoing transcription is inhibited to facilitate accurate DNA repair while transcriptional recovery occurs after DNA repair is complete. However, the mechanisms at play and identity of the transcripts being regulated in this manner are unclear. In contrast to the situation following UV damage, we found that transcriptional recovery after ionizing radiation (IR) occurs in a manner independent of the HIRA histone chaperone. Sequencing of the nascent transcripts identified a programmed transcriptional response, where certain transcripts and pathways are rapidly downregulated after IR, while other transcripts and pathways are upregulated. Specifically, most of the loss of nascent transcripts occurring after IR is due to inhibition of transcriptional initiation of the highly transcribed histone genes and the rDNA. To identify factors responsible for transcriptional inhibition after IR in an unbiased manner, we performed a whole genome gRNA library CRISPR / Cas9 screen. Many of the top hits in our screen were factors required for protein neddylation. However, at short times after inhibition of neddylation, transcriptional inhibition still occurred after IR, even though neddylation was effectively inhibited. Persistent inhibition of neddylation blocked transcriptional inhibition after IR, and it also leads to cell cycle arrest. Indeed, we uncovered that many inhibitors and conditions that lead to cell cycle arrest in G_1_ or G_2_ phase also prevent transcriptional inhibition after IR. As such, it appears that transcriptional inhibition after IR occurs preferentially at highly expressed genes in cycling cells.

## Introduction

DNA double strand breaks (DSB) are one of most deleterious types of DNA lesions. Failure to repair a single DSB can lead to loss of a chromosome arm or cell death and inaccurate repair can lead to changes such as insertions, deletions, and translocations. Accordingly, the cell has developed an intricate DNA damage response (Jackson and Bartek, 2009). In vertebrate cells, the DNA damage response is mediated through activation of three PI3-like kinases: ataxia telangiectasia mutated (ATM), Ataxia telangiectasia and Rad3-related (ATR) and DNA dependent protein kinase (DNA-PK) (Blackford and Jackson, 2017), which coordinates DNA repair and the DNA damage cell cycle checkpoint which arrests cells until DSBs are repaired.

Additionally, ATM and DNA-PK have been shown to inhibit transcription in response to DSBs in a manner that occurs so transiently that it can only be detected when examining the nascent, not bulk, transcripts (Pankotai and Soutoglou, 2013). The transcription resumes or “recovers” immediately after DSB repair (Pankotai and Soutoglou, 2013). This transient inhibition of transcription after DSBs was initially shown for RNA polymerase I (Pol I) transcripts, where ATM triggered the reduction of nascent ribosomal gene transcripts, shown by visualization of a labelled ribonucleotide analog within the nucleolus, after exposure to ionizing radiation (IR) (Kruhlak et al., 2007). Mechanistically, the DSBs triggered a reduction in Pol I initiation complex assembly and led to premature displacement of elongating Pol I from the rDNA genes (Kruhlak et al., 2007). Using a reporter that allowed for visualization of repair protein recruitment and local transcription within cells, it was subsequently shown that ATM also mediates the inhibition of RNA polymerase II (Pol II) transcriptional elongation of genes in the vicinity of I-SceI endonuclease induced DSBs (Shanbhag et al., 2010). This transcriptional inhibition was partly dependent on the E3 ubiquitin ligases RNF8 and RNF168, whereas transcriptional recovery depended on the USP16 enzyme that deubiquitylates histone H2A (Shanbhag et al., 2010). Additional mechanistic analyses using this same system revealed that ATM dependent phosphorylation of the ATP-dependent nucleosome remodeler PBAF is required for local transcriptional inhibition of Pol II transcription flanking a DSB (Kakarougkas et al., 2014), indicating that chromatin changes are also required for transcriptional inhibition in response to DSBs. The purpose of local transcriptional inhibition is to allow efficient and accurate DSB repair (Kakarougkas et al., 2014, Meisenberg et al., 2019). Polycomb group proteins and cohesin have also been shown to be required for local transcriptional inhibition of Pol II transcription flanking a DSB, although their role is unclear (Meisenberg et al., 2019, Kakarougkas et al., 2014), further indicating that chromatin structure and potentially chromosome architecture also regulate transcriptional inhibition in response to DSBs.

Somewhat surprisingly, a distinct mechanism has been reported for the inhibition of Pol II transcription at genes containing a DSB induced by the endonuclease I-PpoI (Pankotai et al., 2012). In this case, both Pol II initiation and elongation were reduced adjacent to the DSB, in a manner dependent on DNA-PK and the proteasome (Pankotai et al., 2012). Mechanistically, DNA-PK appeared to help recruit the E3 ubiquitin ligase WWP2 to DSBs, which then promoted the proteosome-dependent eviction of Pol II (Caron et al., 2019). In the absence of WWP2, the DNA repair machinery was not efficiently recruited, indicating again that the reason for transcriptional inhibition *in cis* flanking a DSB is to promote DNA repair (Caron et al., 2019). The papers examining transcriptional inhibition around DSBs induced by endonucleases generally find the transient repression occurs locally or *in cis* to the DSB (Iannelli et al., 2017). However, another study found induction of the same set of ∼ 200 transcripts soon after irradiation and endonuclease break induction that occurred in a manner dependent on ATM and p53, while only 33 nascent transcripts were down regulated after DSB induction (Venkata Narayanan et al., 2017). Yet another study found that more genes were repressed than induced after inducing global DSBs with neocarzinostatin, and this occurred via p53 mediated down-regulation of MYC (Porter et al., 2017). As such, there are contradictory findings in the field at present. Furthermore, the mechanism of transcriptional recovery after DSB repair is far from clear. In response to UV damage, global transcriptional inhibition and recovery occurs, and this transcriptional recovery after UV repair is dependent on the histone variant H3.3 histone chaperone HIRA (Bouvier et al., 2021, AdamPolo and Almouzni, 2013). Mechanistically, HIRA functioned to repress the transcriptional repressor ATF3, in turn promoting transcriptional recovery after UV repair (Bouvier et al., 2021).

In contrast to the studies to date that have only examined local transcription inhibition occurring *in cis* after DSB damage, we sought to examine transient transcriptional inhibition after induction of global DSBs by IR exposure, and the subsequent transcriptional recovery after DSB repair, via fluorescent labelling of a ribonucleotide analog incorporated only into nascent transcripts. Unlike the situation following UV repair, we do not find a role for HIRA in transcriptional recovery after DSB repair. Our sequencing of the nascent transcripts after irradiation identified a programmed transcriptional program where a larger number of protein-coding genes were upregulated than downregulated. The genes that were immediately downregulated after IR tended to be highly transcribed genes including the rRNAs and histones, while the upregulated genes tended to be transcribed at a lower level. We developed a flow cytometry-based assay of nascent transcripts and used it as the basis for a whole genome gRNA screen to identify factors required for transcriptional inhibition after IR. In addition to finding ATM as being required for inhibition of transcription after DSB induction, we found that depletion of factors leading to cell cycle arrest also blocked transcriptional inhibition.

## Results

### HIRA independent transcriptional inhibition and recovery after ionizing radiation

To detect bulk changes in nascent transcripts after irradiation *in situ*, we added the uridine analog ethynyl uridine (EU) to human U2OS cells for 30 minutes. The incorporated EU was detected by click chemistry to a fluorescent azide followed by immunofluorescence microscopy (Jao and Salic, 2008). A reduction in bulk nascent transcripts was apparent 30 minutes after exposure to 10 Gray ionizing radiation (IR), and the transcriptional recovery was already occurring two hours after irradiation (Fig. 1A). Co-immunofluorescence with γH2AX showed that the DNA damage signal was greatly reduced at the same time point after IR where transcriptional recovery occurred (Figure 1-figure supplement 1), consistent with transcriptional recovery occurring after DSB repair. Given that the histone variant H3.3 histone chaperone HIRA promotes transcriptional recovery after UV repair (Bouvier et al., 2021, AdamPolo and Almouzni, 2013), we tested whether that was also the case for transcriptional recovery after IR. We found that shRNA depletion of HIRA (Figure 1-figure supplement 2) had no effect on transcriptional inhibition nor recovery after IR (Fig. 1A). In agreement, depletion of transcripts encoded from both H3.3 genes (Figure 1-figure supplement 2) had no effect on transcriptional inhibition or transcriptional recovery after IR (Fig. 1B). As such, the requirement for HIRA for transcriptional recovery differs following IR and UV exposure, suggesting differences in the mechanism of these processes.

**Figure 1.**
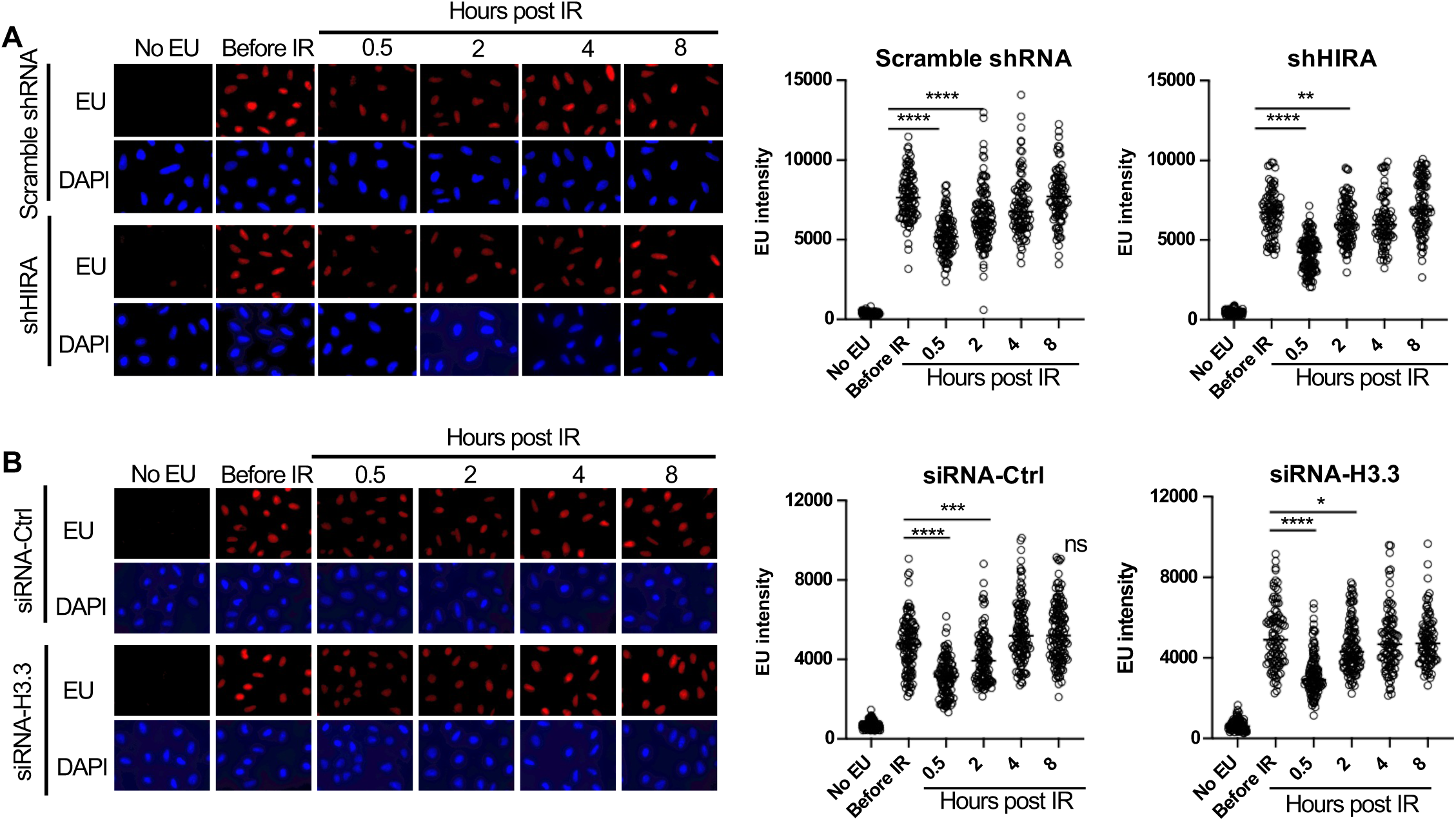
Transcriptional inhibition after irradiation and transcriptional restart after DNA repair in a HIRA independent manner. **A.** U2OS cells were transfected with either a scrambled shRNA or shRNA against HIRA, and were either incubated with EU or not, as indicated, and were irradiated (10 Gy) or not as indicated, followed by detection of EU by click chemistry of a fluorophore and DNA was detected by DAPI staining. The right panel shows quantitation of the mean intensity of EU in at least 80 cells for each condition. **** indicates p<0.001, ** indicates p<0.01, by students T-test. **B.** U2OS cells were transfected with either a control siRNA (siRNA-Ctrl) or two siRNAs against each gene encoding H3.3 (siRNA-H3). EU and DAPI were detected as described in A and quantitated as described in A. **** indicates p<0.001, *** indicates p<0.005, * indicates p<0.05 by students T-test.

### Establishment of a flow cytometry-based assay for transcriptional inhibition and recovery after irradiation

Given that the read out of nascent EU labelled transcripts after irradiation is fluorescence, we established a flow cytometry-based assay to allow us to screen for factors regulating transcriptional inhibition and recovery after IR. We established this assay in a murine Abelson virus transformed pre-B cell line (termed Abl pre-B cells) (Bredemeyer et al., 2006) which are non-adherent and are stably transformed with doxycycline inducible Cas9 (Chen et al., 2021). By flow cytometry analysis, effective incorporation of EU into nascent transcripts was apparent in a manner dependent on ongoing transcription because it was inhibited by the global RNA polymerase inhibitor Actinomycin D (Fig. 2A). The EU signal had 2 peaks (Fig. 2A) and we asked whether this reflects cell cycle differences in the cells with higher and lower EU incorporation into the nascent transcripts. We conducted cell cycle analysis by labeling newly synthesized DNA with BrdU and staining DNA contents with FxCycle Violet at the same time as using EU to label nascent transcripts. It was apparent that the peak with less EU was from the G_1_ phase cells, while the peak with more EU was derived from S and G_2_ phase cells (Fig. 2B). To determine whether there was a detectable reduction in EU incorporation by flow cytometry after irradiation, we irradiated cells, waited different lengths of times before EU labelling of nascent RNA (Fig. 2C). The transcriptional inhibition after irradiation was clearly detectable by flow cytometry as early as 15 minutes after IR, while transcriptional recovery was complete by 4 hours after IR (Fig. 2D). Also, the extent of transcriptional repression was similar regardless of whether the IR dose was 2, 5 or 10 Gray (Fig. 2D, Figure 2-figure supplement 1). This is consistent with the possibility that the reduction of nascent transcripts after IR is a programmed / signaling response rather than due to proximity of the genes to the DSB, which would have led to a dose response.

**Figure 2.**
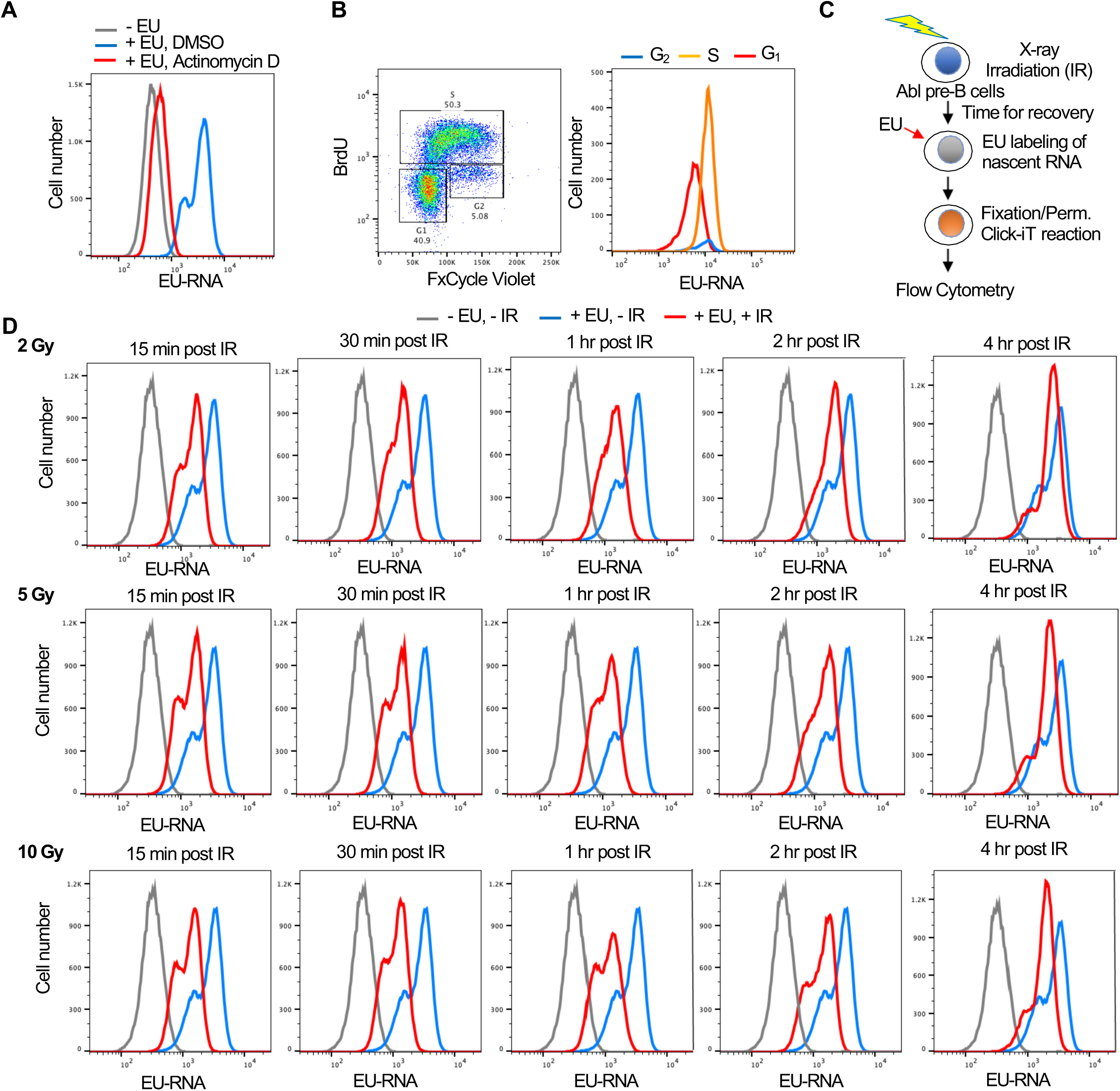
Development of a flow cytometry assay for nascent transcripts shows that transcriptional inhibition after IR is not dose dependent. **A.** The EU positive signal in murine Abl pre-B cells detected by flow cytometry is due to transcripts, as indicated by the addition of 5µM of the general RNA polymerase inhibitor Actinomycin D for 1 hour. B. The two EU peaks observed by flow cytometry correspond to G_1_ (low peak) and G_2_ (high peak) cells. Cycling Abl pre-B cells (left panel) were gated for those with 2N DNA (G_1_) content or 4N (G_2_) content as detected by FxCycle Violate or with BrdU incorporation (S) as indicated and were individually analyzed for EU incorporation into nascent transcripts (right panel). C. Schematic of the assay to detect transcriptional inhibition and transcriptional recovery after IR. D. Time course of transcriptional inhibition and recovery in Abl pre-B cells after IR with the indicated times after IR at the indicated doses of IR.

### Bulk reduction of nascent transcripts after IR is mainly due to a decrease in rDNA and histone gene transcription

To gain a better understanding of transcriptional inhibition after IR, we sought to identify the genes whose transcription was being inhibited after IR. We isolated EU labelled nascent total RNA transcripts 30 minutes after IR and prior to IR from two independent experiments (Figure 3-figure supplement 1) and sequenced the EU-RNA (Fig. 3A). Prior to isolation of the ER labelled nascent RNA, we added equal amounts of the commercial ERCC spike-in to RNA from the same number of cells. This enabled the subsequent normalization of the total read number from the human genome to total reads from the ERCC control, to detect global changes between samples (Chen et al., 2015). We observed that the total read count of nascent transcripts declined after IR (Fig. 3B). Most of the read counts were due to rDNA transcripts, and the decline in bulk transcripts after IR was mostly due to a significant decline in rDNA transcripts (Fig. 3B, Figure 3-figure supplement 2A). By contrast, the total read count from the protein coding transcripts significantly increased after IR (Fig. 3B). Analysis of the protein coding transcripts showed that the transcripts of 3,026 and 1,388 protein-coding genes increased and decreased after IR, respectively (Fig. 3C and 3D, Supp. File 1). To validate our EU-RNA sequencing results, we performed quantitative RT-PCR to measure the nascent transcript levels of genes that were up- and down-regulated after IR. Consistently, we found rDNA transcripts of *28S* and *18S* were significantly downregulated after IR; while the p53 regulated gene *Cyclin-dependent kinase inhibitor 1* (*Cdkn1a*/p21) was highly induced (Figure 3-figure supplement 3). The gene ontology terms describing the genes that were activated after IR included known DNA damage response pathways and related genes (Fig. 3E, Supp. File 2). For examples, the intrinsic apoptotic signaling pathway in response to DNA damage (GO:0008630), type 2 response (GO:0042092), and cytokine-mediated signaling pathway (GO:0019221) were up regulated significantly. Pro-inflammatory cytokines are the major components of immediate early gene programs, being rapidly activated after irradiation in various cell types (SchaueKachikwu and McBride, 2012) ultimately leading to radiation-induced fibrosis in cancer patients following radiation therapy (KimJenrow and Brown, 2014) (Yu et al., 2023). Meanwhile many of the genes that were downregulated after IR included gene products that are involved in chromatin organization and nucleosome assembly (Fig. 3E, Supp. File 3). For examples, chromatin silencing (GO:0006342), DNA packaging (GO:0006323), and nucleosome assembly (GO:0006334) genes were down regulated significantly.

**Figure 3.**
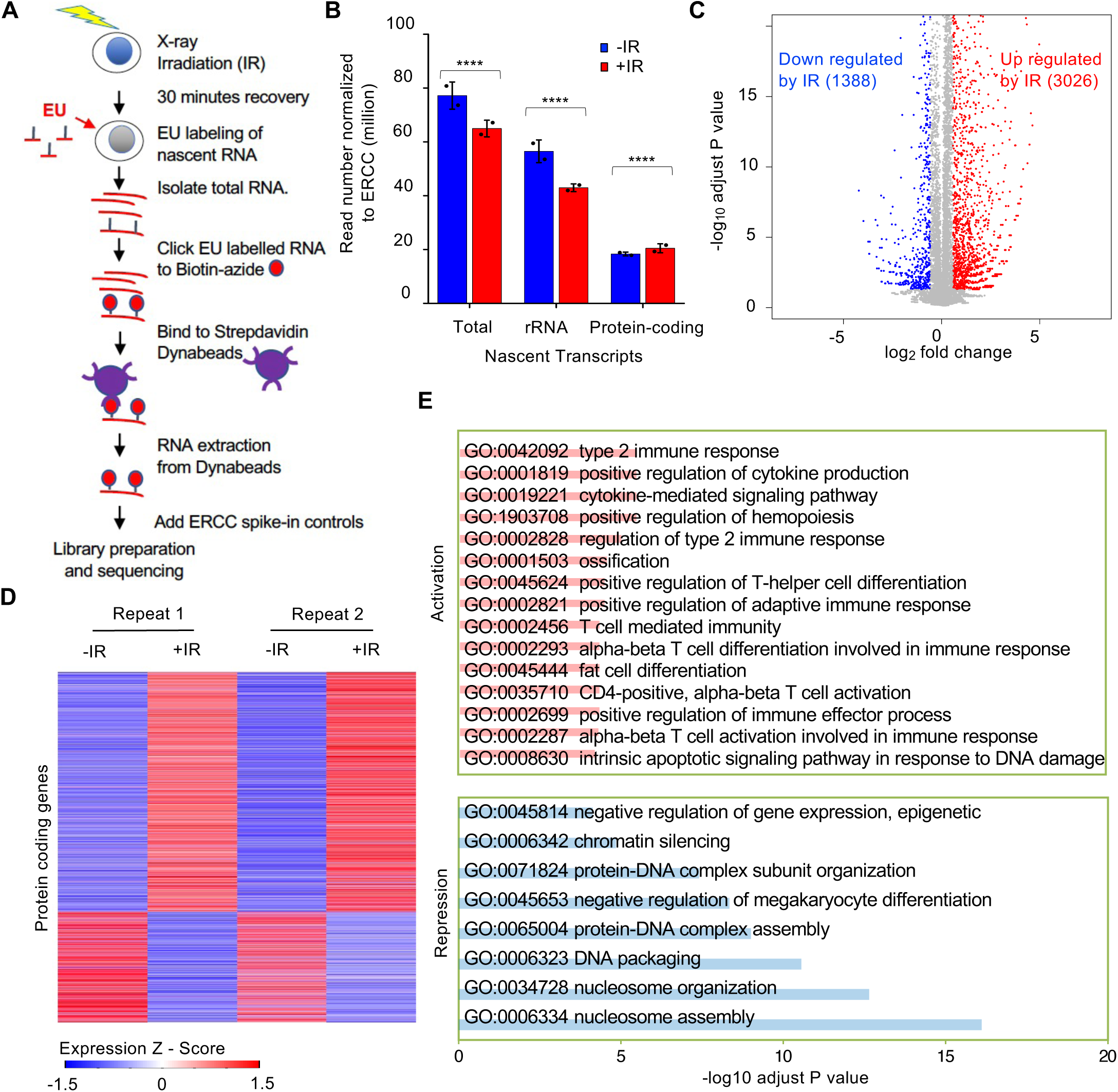
Reduction of nascent transcript levels after irradiation is mainly of the rDNA. **A.** Schematic of nascent transcript sequencing. **B.** Read counts for the total nascent transcripts, rDNA transcripts and protein coding transcripts before and 30 minutes after IR, normalized to ERCC spike in controls. **C**. Significantly changed nascent transcripts from protein coding genes are indicated upon irradiation, and the numbers indicate the number of upregulated and downregulated genes 30 minutes after IR. Data shown are an average of the two independent experimental repeats. **D.** Heat map of significantly increased and decreased nascent transcripts 30 minutes after IR, shown for two independent experimental repeats. Expression z-score was calculated by subtracting the overall average gene abundance from the raw expression for each gene and dividing that result by the standard deviation (SD) of all of the measured counts across all samples. **E.** Gene Ontology analysis of the top significantly enriched GO terms most upregulated after IR (pink) and most downregulated after IR (blue). Enriched gene number (red) and fold enrichment (blue) were showed in each GO term.

To gain more insights on how the transcription of protein coding genes was regulated after IR, we defined differentially expressed genes (DEGs) between samples before and after IR. We found read number from these DEGs became significantly greater after IR (Fig. 4A). We sorted the protein-coding DEGs by average expression level of each gene in all 4 samples: two replicate samples before IR and two replicates after IR. The number of reads derived from the mostly highly expressed protein-coding DEGs became significantly smaller after IR (Fig. 4A, 4B). If the gene repression after IR is due to their being *in cis* to the DNA lesion, it would be expected that genes that were repressed after IR would tend to be longer, because they would be more likely to be damaged. However, this was not the case because the length of the nascent transcripts was equivalent regardless of whether their transcription was repressed, activated, or not changed after IR (Fig. 4C). Intriguingly, we found that the repressed genes of the top 100 high-expression DEGs tended to be shorter (Fig. 4C). Next, we inspected the expression level of individual protein-coding genes and confirmed that most changes in gene expression after IR tended to occur for the genes that were activated after IR, while many of the genes that had a high-expression level were repressed after IR, for example, the histone encoding genes (Fig. 4D). Strikingly, we found that vast majority of the histone genes showed reduced transcription after IR (Fig. 4E, Figure 3-figure supplement 2B), and was validated by RT-PCR analysis (Figure 3-figure supplement 3). Finally, to determine whether transcriptional repression was occurring at the initiation, elongation or both stages of transcription, we examined the read counts throughout the open reading frames of the repressed protein coding genes, before and after IR. We found that the decrease of transcripts mainly occurred in the gene body of these genes with similar intensity at both the 3’ and 5’ ends of the gene body, which indicates transcriptional repression after IR occurred at the initiation stage of transcription (Fig. 4F). Therefore, these data indicate that the bulk reduction in nascent transcripts after IR is mainly due to reduced transcriptional initiation of the rDNA and histone genes.

**Figure 4.**
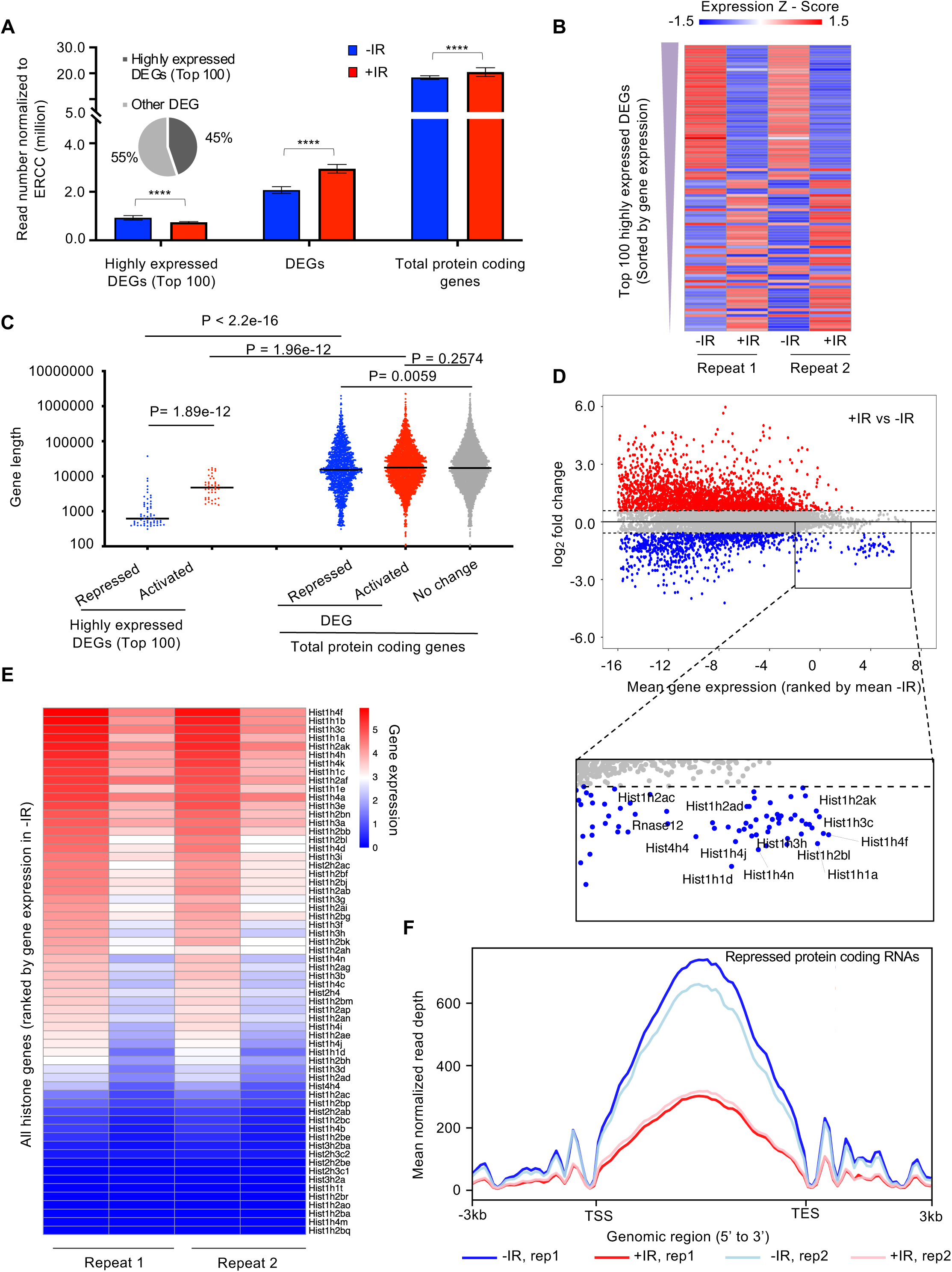
The highly transcribed protein coding genes tend to be repressed after irradiation, due to a decrease in the transcription of the histone genes. **A.** Plot of the transcript abundance of differentially expressed genes (DEGs) showing that highly expressed genes have reduced nascent transcript levels 30 minutes after IR, while moderately expressed and low level expressed genes tend to have increased nascent transcript levels 30 minutes after IR. Mean gene expression and standard deviation is shown in million reads mapped to genes normalized by ERCC spike-in reads. Data are shown from two independent experimental repeats (rep) of the experiment. B. Heat map showing nascent transcript levels of the top 100 highly expressed DEGs, ranked by gene expression from top (highest) to bottom, 30 minutes after IR, shown for two independent experimental repeats. Expression z-score was calculated by subtracting the overall average gene abundance from the raw expression for each gene and dividing that result by the standard deviation (SD) of all the measured counts across all four samples. C. Among the top 100 of highly expressed protein-coding genes, repressed genes are significantly shorter compared to activated genes. The activated, non-changed and repressed genes show little difference in gene size (the data are averaged for each gene between two independent experimental repeats). D. Plot of change in gene expression after IR against mean gene expression (log_2_), ranked by mean gene expression in samples before IR on the x axis, for all nascent transcripts. Some of the highly expressed genes whose nascent transcript levels decreased after IR are labelled in the rectangle, including histone genes. E. Heat map of nascent transcripts of all histone genes 30 minutes after IR, shown for two independent experimental repeats. F. The average read counts for repressed protein coding genes throughout their gene length before and after IR for 2 independent repeats of the experiment.

### A CRISPR-Cas9 screen identifies ATM, neddylation and CUL4B as promoting transcriptional inhibition after IR

We performed a genome-wide gRNA library CRISPR-Cas9 screen in Abl pre-B cells, allowing 7 days for the gRNAs to inactivate their target genes (Fig. 5A). We then sorted the 10% of the cells with the most nascent RNA (high EU) 30 minutes after IR, as these would include cells with gRNAs corresponding to gene products that are required for transcriptional inhibition after IR. We sequenced the gRNAs within the high EU cells and within the total cell population. We calculated an enrichment score for each of the 5 gRNAs that were included in the library against each gene (Supp. File 4, Figure 5-figure supplement 1). Fig. 5B shows the enrichment scores for the 5 gRNAs for some of the top hits (high EU) from the screen. We found that gRNAs against ATM were enriched in the high EU cells, consistent with the fact that transcriptional inhibition of a gene after induction of a DSB requires ATM signaling (Shanbhag et al., 2010). Other top hits included most of the machinery that mediates neddylation, the post translational covalent addition of NEDD8, a small ubiquitin-like peptide, onto other proteins (Rabut and Peter, 2008). These hits included the *Nedd8* gene encoding the NEDD8 ubiquitin like modifier, *Nae1* encoding NEDD8 activating enzyme E1 subunit 1 NAE1, *Uba3* encoding the Ubiquitin like modifier activating enzyme 3 UBA3, *Ube2m/Ubc12* encoding the NEDD8-conjugating enzyme UBC12, *Ube2f* encoding a neddylation E2 enzyme UBE2F, and *Rbx1* and *Rbx2* which encode linkers that facilitate NEDD8 transfer from the E2 enzyme to Cullins (Fig. 5B). We also uncovered gRNAs against the gene encoding the neddylation substrate CUL4B enriched in the high EC cells (Fig. 5B). These data suggested that neddylation may have a novel role in transcriptional inhibition after IR.

**Figure 5.**
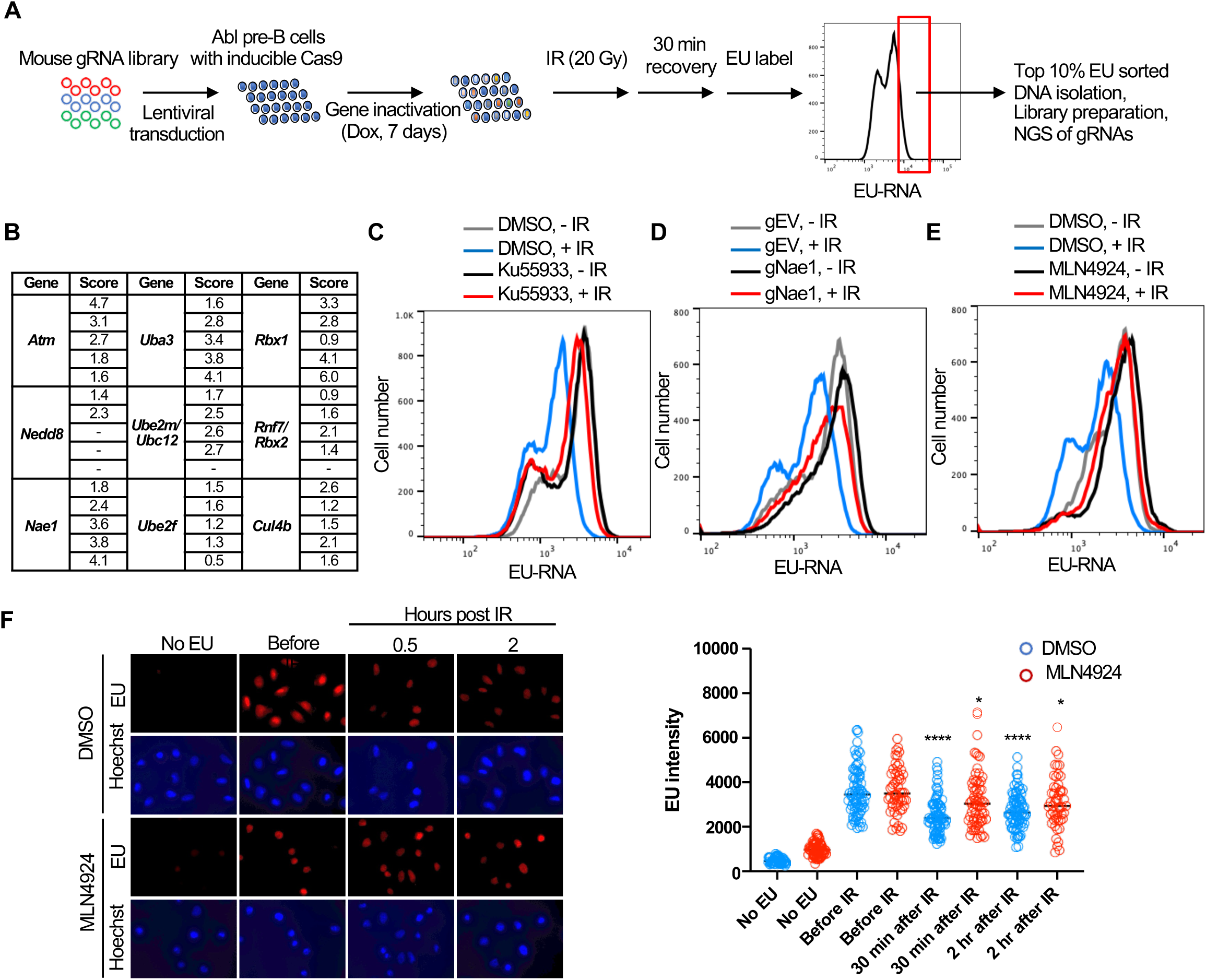
Whole genome gRNA screen CRISPR-Cas9 screen for factors involved in transcriptional inhibition after irradiation identifies the neddylation pathway. **A.** Schematic of whole genome gRNA CRISPR-Cas9 screen for gene products that promote transcriptional inhibition after IR. B. Fold enrichment of 5 guide RNAs against the indicated genes in the 10% of cells with most EU incorporated into transcripts 30 minutes after IR. C. Inhibition of ATM greatly reduces transcriptional inhibition 30 minutes after IR in Abl pre-B cells. The inhibitor was used at 15 µM for 1 hours. D. gRNA mediated depletion of Nae1 greatly reduces transcriptional inhibition 30 minutes after IR in Abl pre-B cells. E. Inhibition of neddylation greatly reduces transcriptional inhibition 30 minutes after IR. The inhibitor was used at 1µM for 16 hours in Abl pre-B cells. F. Inhibition of neddylation reduces transcriptional inhibition after IR in U2OS cells, as detected by fluorescence analysis of nascent transcripts as described in legend to Fig. 1 and quantitated as in Fig. 1. Significant difference after IR compared to before IR are indicated by asterisks, where **** indicates p<0.001, * indicates p<0.05 by students T-test. All experiments in this figure are in murine Abl pre-B cells.

We validated the role of ATM in bulk transcriptional inhibition after IR with the ATM inhibitor Ku55933 (Fig. 5C). To validate a role for neddylation in transcriptional shut off after IR, we depleted NAE1 with a gRNA to *Nae1* (Figure 5-figure supplement 2) and showed that depletion of NAE1 also greatly reduced transcriptional inhibition after IR (Fig. 5D). Similarly, a 16-hour treatment with the neddylation inhibitor MLN4924 also greatly reduced transcriptional inhibition after IR (Fig. 5E). To ensure that the role of neddylation in transcriptional inhibition after IR is not unique to the murine Abl pre-B cells, we found that transcriptional inhibition, detected by fluorescence microscopy, in U2OS cells was significantly reduced upon inhibition of neddylation (Fig. 5F). As such, these data suggest that neddylation is required for efficient transcriptional inhibition after IR.

While most neddylation substrates have only been identified upon overexpression of the neddylation machinery, the Cullins have been shown to be *bone fide* neddylation substrates *in vivo* (Pan et al., 2004). Given that we found gRNAs to Cul4b encoding CULLIN 4B enriched in the EU high cells (Fig. 5B) we asked whether its neddylation is induced after IR. We found that when compared to the level of unneddylated CUL4A there was not a significantly higher proportion of neddylated CUL4A or CUL4B in U2OS cells after IR (Fig. 6A). We used the neddylation inhibitor MLN4924 as a positive control to confirm which bands were neddylated CUL4A and CUL4B (Fig. 6A). To determine whether CUL4B or CUL4A were required for IR induced transcriptional inhibition, we made stable Abl pre-B cell lines lacking each protein by gRNA-mediated disruption of the *Cul4a* or *Cul4b* genes. Loss of CUL4A had no effect on transcriptional inhibition after IR, while loss of CUL4B partially reduced transcriptional inhibition after IR (Fig. 6B, Figure 6-figure supplement 1A). Given that CUL4B and CUL4A show some functional redundancy (Hannah and Zhou, 2015, Brown et al., 2015), we depleted both proteins at the same time. Additional transient bulk gRNA transfection-mediated depletion of CUL4A from cells lacking CUL4B (depletion of both CUL4A and CUL4B is lethal) did not further increase the block of transcriptional inhibition after IR (Figure 6-figure supplement 1B). As such, these data indicate that CUL4B but not CUL4A contributes to transcriptional inhibition after IR. Given that neddylation of CUL4A/CUL4B regulates cell cycle progression (Hannah and Zhou, 2015), and we saw changes in the distribution of the cell cycle phases upon CUL4A/CUL4B depletion and upon blocking neddylation as indicated by the change in relative heights of the EU low (G_1_) and EU high (S/G_2_) peaks (Fig. 5D, 5E, 6B and Figure 6-figure supplement 1B,C), we wondered if cell cycle arrest may be influencing transcriptional inhibition after IR. Accordingly, we inhibited neddylation for 1-3 hours in Abl pre-B cells, which effectively inhibited neddylation (Fig. 6C) and tested the effect on transcriptional inhibition after IR. Strikingly, we observed transcriptional inhibition after IR even upon neddylation inhibition treatment for 1-3 hours (Fig. 6D), indicating that neddylation is not required for transcriptional inhibition after IR. We also observed that the length of time required for treatment with the neddylation inhibition to block transcriptional inhibition after IR (Fig. 5E) caused arrest of the cell cycle in G_2_ phase (Figure 6-figure supplement 2). Therefore, these data indicate that neddylation *per se* is not required for transcriptional inhibition after IR. They also raise the possibility that the cell cycle arrest caused by persistent loss of neddylation may prevent transcriptional inhibition after IR.

**Figure 6.**
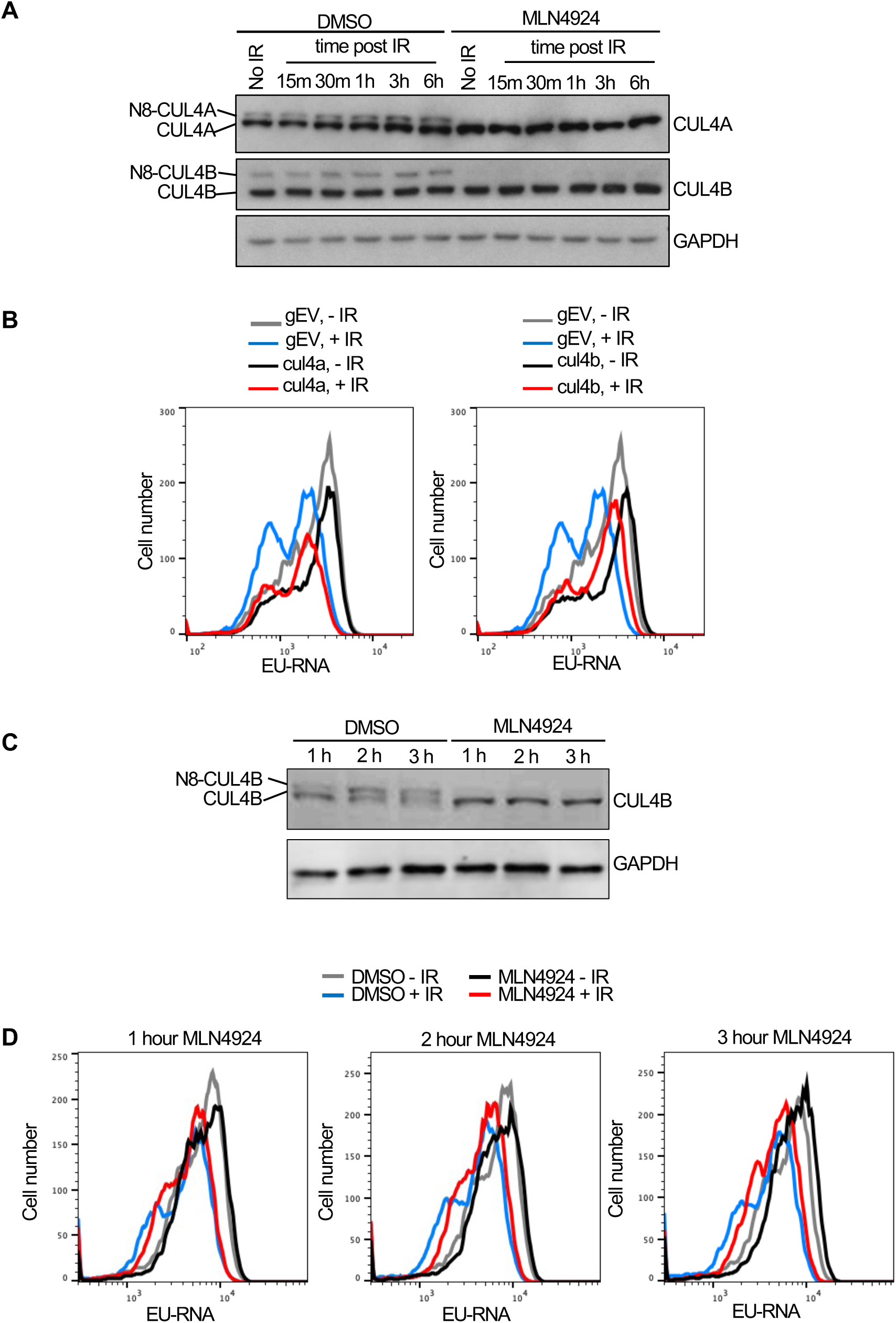
CUL4B but not CUL4A contributes to transcriptional inhibition after irradiation. **A.** Analysis of CUL4A and CUL4B neddylation in U2OS cells after IR, in the absence of presence of 10 µM treatment for 3 hours with the neddylation inhibitor. N8 indicates the neddylated species. **B.** Analysis of nascent transcripts in Abl pre-B cell lines stably depleted of CUL4A or CUL4B 30 minutes after IR, as indicated. **C.** Short treatment of Abl pre-B cells with neddylation inhibitor MLN4924 is sufficient to block neddylation of CUL4B. **D.** The same cells used in C were analyzed for EU incorporation into nascent transcripts 30 minutes after irradiation, without or with the indicated time of MLN4924 treatment before irradiation.

### Cell cycle arrest in G_1_ or G_2_ phase blocks inhibition of transcription after IR

To directly determine the relationship between cell cycle arrest caused by prolonged neddylation inhibition and transcriptional inhibition after IR, we labelled the Abl pre-B cells with both EU (nascent transcripts) and 7-Aminoactinomycin D (7-AAD) (DNA stain). Prolonged neddylation inhibition led to accumulation of cells with a 4N DNA content and greatly reduced transcriptional inhibition after IR (Fig. 7A, Figure 6-figure supplement 2). To investigate if this correlation was specific to neddylation inhibition, we used an unrelated inhibitor that causes cell cycle arrest with a 4N DNA content, RO-3306, a CDK1 inhibitor (Vassilev et al., 2006). We also saw accumulation of cells with a 4N DNA content and no transcriptional inhibition after IR upon CDK1 inhibition (Fig. 7B, Figure 6-figure supplement 2). To determine whether this effect was unique to cells arrested with a 4N DNA content or was shared with conditions that cause arrest in G_1_ phase, we treated cells with 10% or 0.1% FBS, where the later lead to arrest in G_1_ phase (due to serum depletion) and prevented transcriptional inhibition after IR (Fig. 7C). It is relevant to point out that the level of bulk nascent transcripts in cells arrested in G_1_ by serum depletion is still far higher than cells lacking EU (Fig. 7C), suggesting that the limited transcriptional inhibition after IR in G_1_ arrested cells is not just because there is only minimal transcription occurring. In agreement, two CDK4/6 inhibitors, Ribociclib and Palbociclib, that lead to G_1_ arrest also prevented transcriptional inhibition after IR (Fig. 7D, 7E). Intriguingly, these data indicate that multiple different treatments that lead to cell cycle arrest in G_1_ or G_2_ *per se* prevent transcriptional inhibition after IR.

**Figure 7.**
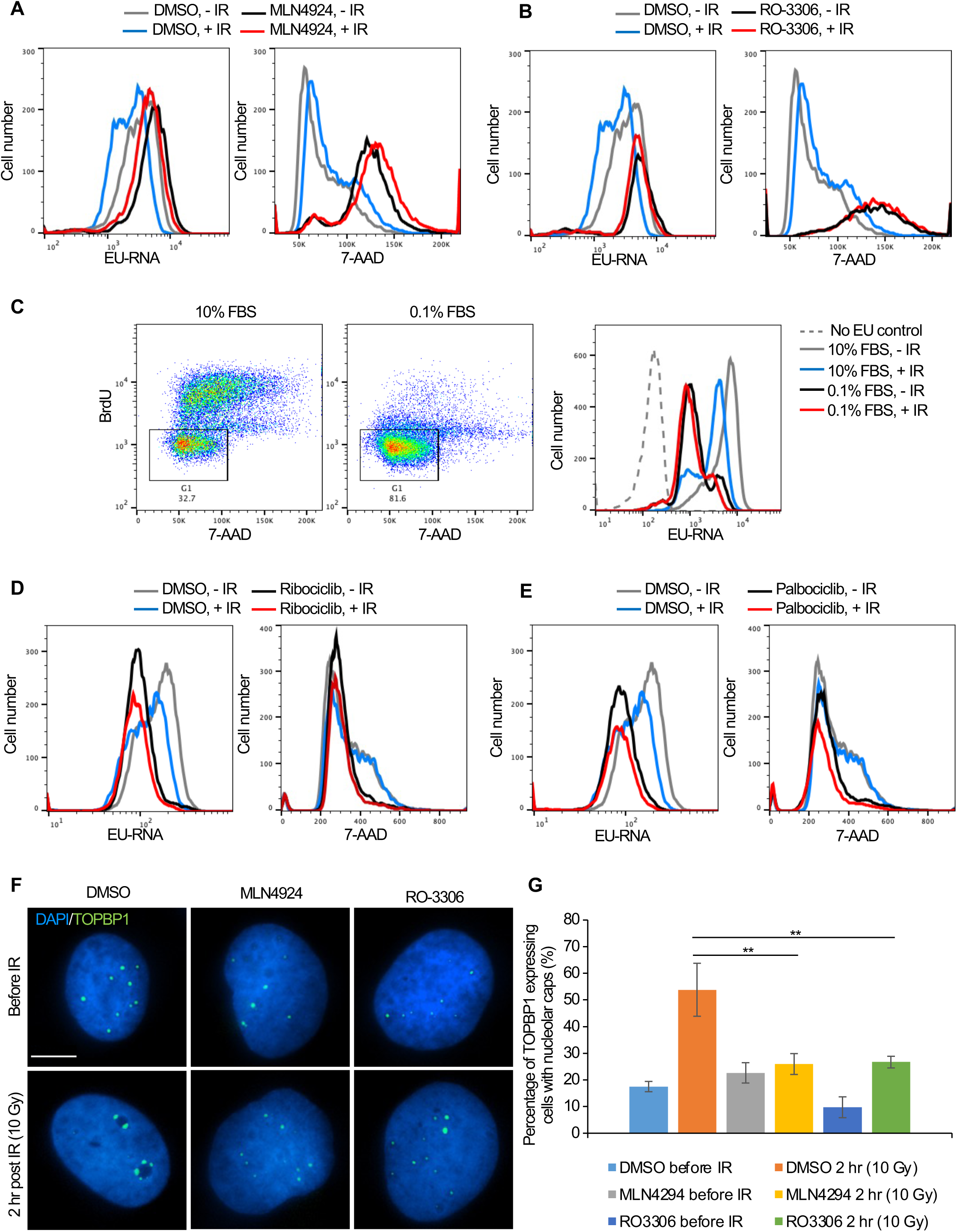
Cell cycle arrest in G_1_ or G_2_ prevents transcriptional inhibition after DNA damage. **A.** Abl pre-B cells were treated with MLN4924 (1 μM) for 16 hours, followed by IR and staining of DNA with 7-AAD and nascent transcripts with EU 30 minutes after IR. **B.** Abl pre-B cells were treated with RO-3306 (10 μM) for 16 hours, followed by IR and staining of DNA with 7-AAD and nascent transcripts with EU 30 minutes after IR. **C.** The left two panels show the cell cycle distribution of Abl pre-B cells after growth for 72 hours in 1% FBS or 0.1% FBS. The rectangles and numbers indicate the % of cells with a 2N DNA content. The right panel shows the EU incorporated into nascent transcripts 30 minutes after IR for the same samples. **D.** Abl pre-B cells were treated with Ribociclib (5 μM) for 24 hr., followed by IR and staining of DNA with 7-AAD and nascent transcripts with EU 30 minutes after IR. **E.** Abl pre-B cells were treated with Palbociclib (5 μM) for 24 hr, followed by IR and staining of DNA with 7-AAD and nascent transcripts with EU 30 minutes after IR. **F.** The U2OS cells were treated with doxycycline (1 μg/mL) for 12 hours to express eGFP-TOPBP1. Then, DMSO, MLN4924 and RO3306 were added to the cells for another 16 hours. TOPBP1 localization was shown in cells before IR or 2 hours after IR (10 Gy). Scale bar is 10 μm. **G.** Quantification of TOPBP1 expressing cells with nucleolar caps before and after IR (10 Gy) in cells treated with MLN4924 and RO-3306. Data shown are an average of the three independent experimental repeats. Significant differences are indicated by asterisks, where ** indicates p<0.01 by students T-test.

To gain molecular insight into why ATM-dependent bulk transcriptional inhibition occurs after IR in cycling cells but not in arrested cells, we investigated the possibility that ATM-dependent pathways that inhibit rDNA transcription after IR in cycling cells (Kruhlak et al., 2007) fail in arrested cells. In cycling cells, DSBs within the rDNA trigger formation of nucleolar caps due to damaged rDNA and associated proteins relocalizing to the nucleolar periphery (van Sluis and McStay, 2017). In cycling cells, DSBs within the rDNA trigger ATM-dependent phosphorylation of Treacle which promotes recruitment of NBS1 and TOPBP1 to the nucleolar caps to inhibit rDNA transcription (Larsen et al., 2014, Mooser et al., 2020). Accordingly, in cycling cells, TOPBP1 accumulates in the nucleolar caps after induction of DSBs, concomitant with repression of rDNA transcription (Sokka et al., 2015, Mooser et al., 2020). We asked whether TOPBP1-eGFP recruitment to nucleolar caps was disrupted after DNA damage in cells arrested by treatment with MLN4924 or RO-3306. The overall induction of TOPBP1 expression was not affected by inhibitor treatment (Figure 7-figure supplement 1). However, the percentage of cells with TOPBP1-eGFP localizing to nucleolar caps after IR was markedly reduced upon MLN4924 and RO-3306 treatment (Fig. 7F, 7G). These results are consistent with arrested cells failing to repress rDNA expression after global DNA damage, as a consequence of compromised ATM-dependent localization of TOPB1 to nucleolar caps.

## Discussion

We find that while the bulk abundance of nascent transcripts is rapidly reduced after IR, more protein coding genes are induced than inhibited after IR. Instead, the reduction in bulk nascent transcript levels that occurs after IR is due to reduced transcriptional initiation of a subset of genes that are the most highly expressed in the cell – the rDNA and histone encoding genes. Notably, bulk transcriptional inhibition after IR did not occur in cells arrested in G_1_ or G_2_ phases of the cell cycle indicating that cells need to be cycling for IR to rapidly inhibit bulk transcription. The length-independent and dose-independent reduction in bulk abundance of nascent transcript after IR (Figs. 5C, Supp. Fig 3) suggests that the reduced bulk abundance of nascent transcripts after IR may occur *in trans* as a programmed event. This is in contrast to studies that have found transcriptional inhibition *in cis* of a gene immediately adjacent to an endonuclease-induced DSB. Our work indicates that the genome wide transcriptional response to DSBs after IR cannot be extrapolated from single gene studies.

Analysis of nascent transcripts at early times after irradiation revealed a different transcriptional response compared to changes in total mRNAs after irradiation (Lieberman et al., 2017). These changes in mRNA levels typically occurred in an IR dose-dependent manner. By contrast, the bulk changes in nascent transcripts occurred in a dose-independent manner (Fig. 2D). Analysis of total mRNAs changes following IR showed altered expression of genes involved in signal transduction, regulation of transcription, and metabolism (Su et al., 2004). Similarly, the upregulated nascent transcript changes identified pathways including signal transduction, while nascent transcripts from protein-coding genes that affected nucleosome assembly and chromatin structure (histones) were downregulated after IR (Fig. 3B, 3E). Finally, most of the gene expression changes detected by analysis of total mRNA continued to increase or decrease over long periods of time, up to 48 hours (Su et al., 2004), whereas the reduction of bulk nascent transcript levels occurred in a very transient manner and was already returning to normal by 4 hours after IR (Fig. 1, Fig. 2D).

How is bulk nascent transcription being inhibited after irradiation? Reminiscent of the ATM dependence on the changes in mRNA levels after IR (Artuso et al., 1995), the reduction in bulk nascent transcript levels after IR was also dependent on ATM (Fig. 5C). Given that most of the reduction in bulk nascent transcript levels was due to the rRNA (Fig. 3B), this is consistent with the previous report of ATM-dependent inhibition of RNA Polymerase I transcription in response to DSBs (Kruhlak et al., 2007). In this case, the Pol I transcription appeared to be inhibited at both the initiation and elongation stages (Kruhlak et al., 2007). Sequencing analysis of nascent transcripts of protein-coding genes whose expression significantly decreased 30 minutes after IR suggested that it is the initiation of Pol II transcription that is inhibited, given that the reduction in sequencing reads at the 5’ end and 3’ ends of open reading frames was equivalent (Fig. 4F). Studies that examined the mechanism of Pol II transcriptional inhibition of one gene adjacent to an endonuclease induced DSB identified a reduction in transcriptional elongation as indicated by reduced Pol II Ser2 phosphorylation (Shanbhag et al., 2010), while other studies found a defect in both Pol II initiation and elongation of a different gene adjacent to an endonuclease induced DSB, in a manner dependent on DNA-PK and the proteasome (Caron et al., 2019, Pankotai et al., 2012). Our data is consistent with the possibility that the major mechanism for the repression of the ∼1,000 protein coding genes after IR is at the transcriptional initiation stage. However, our data do not rule out that transcriptional elongation may be additionally repressed after IR, but would not be observed in our analyses due to the repression of transcriptional initiation.

Rapid inhibition of transcription *in cis* has been observed following endonuclease-mediated DSB induction, where a DSB-proximal transcriptional reporter was inhibited while a second transcriptional reporter inserted elsewhere in the genome without a proximal FokI nuclease site was not inhibited after induction of Fok1 (Shanbhag et al., 2010). If the transcriptional inhibition after IR that we observe was occurring *in cis,* we would expect that longer genes would be more inhibited than shorter genes after IR, as they are more likely to experience a DSB induced by IR. However, this was not the case because the length of the nascent transcripts was equivalent regardless of whether their levels were increased, didn’t change, or were decreased after IR (Fig. 4C). We would also expect that bulk transcript levels would be more reduced with a higher dose of IR if it occurred *in cis*, but that was not the case (Fig. 2D). In fact, transcriptional repression around a nuclease-induced DSB can spread hundreds of kb away from the break, throughout a whole topological associated domain marked by gamma H2AX (Purman et al., 2019). Importantly, our data do not contradict that DSBs can induce transcriptional inhibition *in cis*, rather it likely reflects the random nature of the DNA damage induced by IR is not sufficient to detect inhibition *in cis*, as every cell will have DNA DSBs at different locations. We also are examining the effect following global DNA damage induced IR versus the effect of induction of a single or limited numbers of DSBs by endonuclease induction. It is noteworthy that the nascent transcript levels of more genes rapidly increased after IR than were reduced (Figs. 3 and 4), such that the situation, at least after IR, is more complex than transcriptional inhibition *in cis* to the DSBs.

The change in transcript levels after irradiation tended to depend on the expression level of the genes before irradiation. Those genes that were normally most highly transcribed were repressed after IR, while genes that were normally expressed at intermediate or low levels tended to be induced after IR (Fig. 4A). The mechanistic reason for this is unclear. Among the genes that were most repressed after IR treatment were many of the histone encoding genes (Fig. 4D, 4E). Histone gene expression has been shown to be reduced after IR previously in a manner dependent on ATM and p53, as seen at the mRNA level 10 hours or more after irradiation (Su et al., 2004). By contrast, reduction in nascent histone transcript levels occurred 30 mins after IR (Fig. 4D, 4E). Interestingly, we observed a tremendous expression induction of *Cdkn1a/p21* gene, which encodes a potent cyclin-dependent kinase inhibitor, after IR (Supp Fig. 6). The histone transcriptional coactivator NPAT induces histone transcription when it is phosphorylated by cyclin E/CDK2 (Zhao et al., 1998, Zhao et al., 2000, Ma et al., 2000). As such, highly elevated levels of Cdkn1a/p21 after IR might inactivate cyclin E/CDK2, leading to hypo-phosphorylation of NPAT and an immediate repression of histone gene expression after IR. It also may be relevant that the loci found to be repressed by bulk IR are highly repetitive gene arrays that tend to form nuclear sub-compartments (nucleoli, histone bodies). As such, their likelihood of repeats being in the vicinity of a repeat with an IR-induced DNA lesion in three-dimensional space is high, which may promote their transcriptional repression after IR in *trans*. Moreover, silencing may spread through the relevant nuclear sub-compartments, consistent with the formation of DNA damage compartments described recently (Arnould et al., 2023).

In response to UV exposure, bulk inhibition of transcription occurs, followed by transcriptional recovery after repair of UV-induced damage, where transcriptional recovery after repair of UV-induced damage is dependent on the histone variant H3.3 histone chaperone HIRA (Bouvier et al., 2021, AdamPolo and Almouzni, 2013). Mechanistically, HIRA functioned to repress the transcriptional repressor ATF3, in turn promoting transcriptional recovery after repair of UV-induced damage (Bouvier et al., 2021). We find that neither HIRA nor H3.3 is required for recovery of bulk nascent transcripts after DSB repair (Fig. 1). Additionally, we find that the genes that have altered levels of nascent transcripts after UV damage (Bouvier et al., 2021) and after IR are quite distinct (data not shown). However, the UV studies were performed on nascent mRNA from a human cell line, while our studies were performed on total nascent RNA from a mouse cell line.

Why didn’t our CRISRP/Cas9 screen of factors responsible for bulk nascent transcript inhibition after IR identify previously reported factors involved in Pol II transcriptional inhibition proximal to an endonuclease-induced DSB? This is likely because most of the reduction in bulk nascent transcript level that we were detecting after IR was due mainly to reduced transcript abundance from the rDNA, rather than Pol II transcripts (Fig. 3B). We did find ATM in the screen (Fig. 5B, 5C), and this is consistent with the fact that ATM is required for rDNA transcriptional inhibition after IR (Kruhlak et al., 2007). Many of the most significant hits from the screen encoded factors that are involved in the neddylation pathway and the neddylation substrate CUL4B (Fig. 5D, 5E, 5F and Fig. 6B). However, short times of neddylation inhibition were sufficient to inhibit neddylation but did not prevent transcriptional inhibition after IR (Fig. 6C, 6D). Longer times of neddylation inhibition did prevent transcriptional inhibition after IR (Fig. 5E, 5F), but also caused cell cycle arrest (Fig. 7A, Supp. Fig. 10). These results lead us to speculate that neddylation is not directly involved in transcriptional inhibition after IR, but that neddylation promotes cell cycle progression, and that it is the cell cycle arrest occurring upon neddylation inhibition that prevents the bulk reduction of nascent transcript levels after IR. Consistent with the idea that cell cycle arrest *per se* may be preventing transcriptional inhibition after IR, we also found that CDK1 inhibition leading to G_2_ arrest, and serum starvation and CDK4/6 inhibition leading to G_1_ arrest, also prevented bulk transcriptional inhibition after IR (Fig. 7).

Why would cell cycle arrest in G_1_ or G_2_ phases of the cell cycle prevent transcriptional repression of rDNA and histone genes after IR? Transcription of rDNA is known to be reduced in non-cycling cells when total transcript levels were measured (Moss and Stefanovsky, 1995, O’Mahony et al., 1992) while transcription of histone genes requires ongoing DNA replication (SittmanGraves and Marzluff, 1983). As such, one possibility for failure to see bulk reduction in nascent transcript abundance after IR in arrested cells may be that rDNA and histone transcription is already reduced in G_1_ and G_2_ arrested cells. However, we found that the total level of bulk nascent transcripts during G_2_ arrest before IR were equivalent to, or more than, the level of bulk nascent transcripts before cell cycle arrest (compare grey (no arrest) to black lines (cell cycle arrest) in Figs. 5C-E, 7A, 7B) which suggest that the levels of rDNA transcripts may not be reduced during G_2_ arrest in our experiments given that most of the nascent transcripts are from the rDNA in the conditions of our experiments (Fig. 7B). It is also possible that a factor that is required for transcriptional inhibition of the rDNA after IR is absent or inactive in arrested cells.

In addition to ATM, NBS1 was previously shown to be required for transcriptional inhibition of rDNA after IR (Kruhlak et al., 2007). The mechanism for this was unclear, but it is tempting to speculate that the requirement of ATM for transcriptional inhibition after IR in cycling cells may be mediated through ATM-dependent phosphorylation of Treacle. In this form, Treacle functions with TOPBP1 to promote recruitment of NBS1 to nucleolar caps to repress rDNA transcription (Larsen et al., 2014, Mooser et al., 2020). That TOPB1 fails to relocalize to nucleolar caps after IR in arrested cells (Fig. 7F, 7G) is consistent with a potential loss of ATM-dependent treacle phosphorylation in arrested cells, which would prevent reduction in rDNA transcription after DNA damage. Future experiments will reveal further insight into the cell cycle dependent control of transcriptional inhibition of highly transcribed genes after DNA damage.

## Supplementary

**Figure 1-figure supplement 1.**
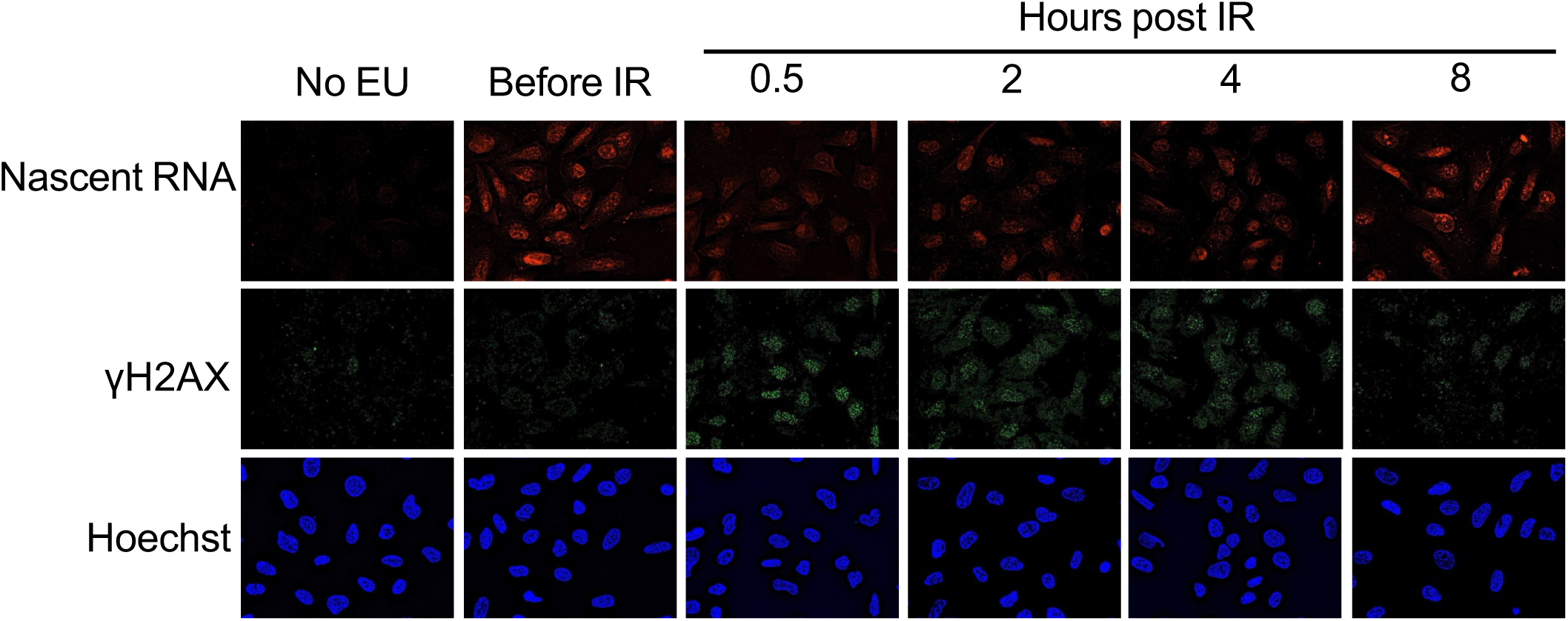
Transcriptional inhibition after irradiation and transcriptional restart after DNA repair in U2OS cells. **A.** U2OS cells were either incubated with EU or not, as indicated, and were irradiated (10 Gy) or not as indicated, followed by detection of the EU by click chemistry of a fluorophore, and immunofluorescence staining of gamma H2AX in the same cells and the DNA was detected by Hoechst staining.

**Figure 1-figure supplement 2.**
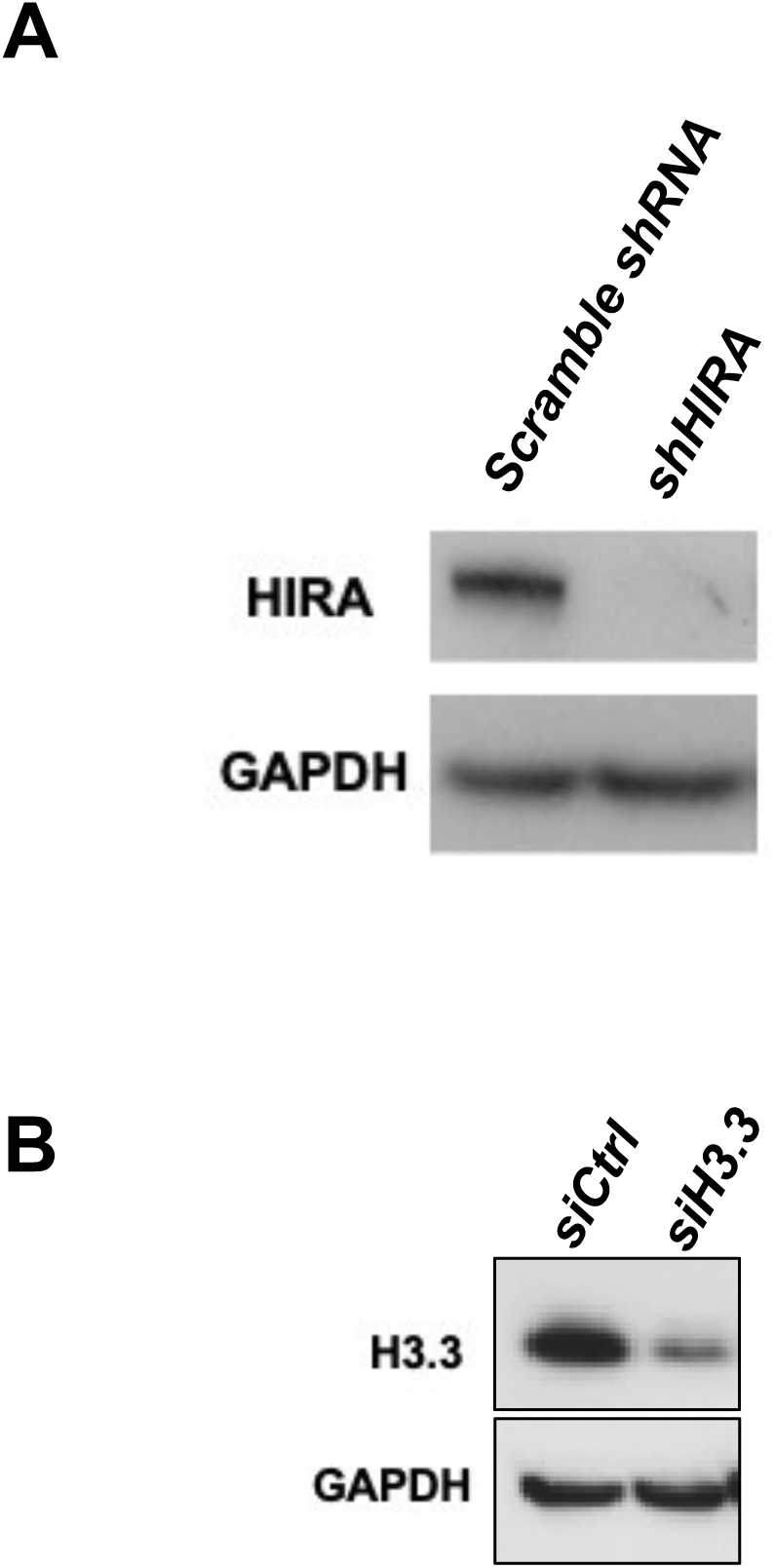
Confirmation of knockdown of HIRA (A) and H3.3 (B). The samples were from the same experiments shown in Figure 1.

**Figure 2-figure supplement 1.**
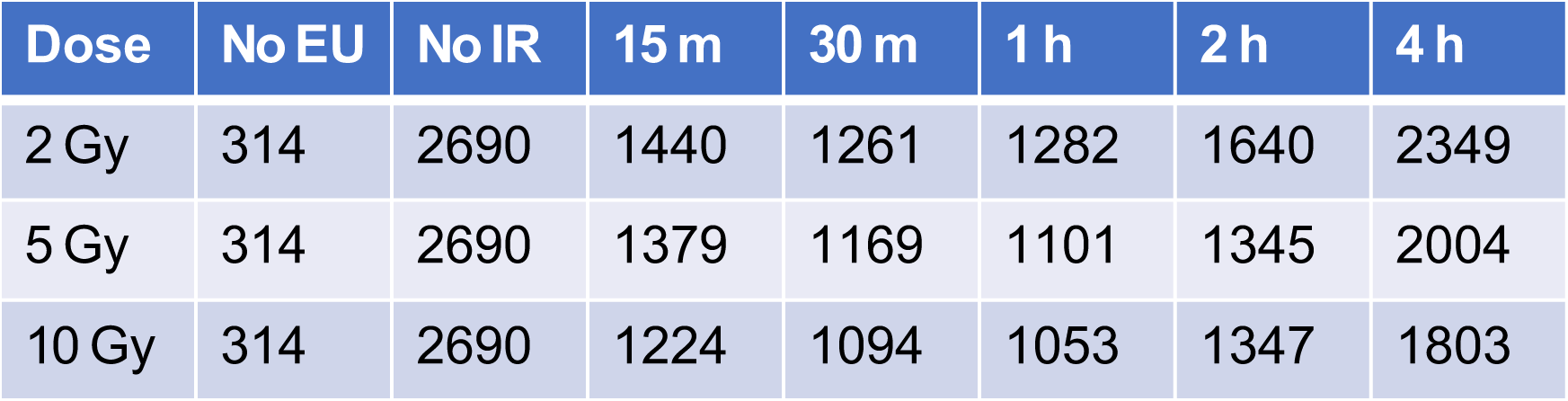
The mean intensities of the EU peaks shown in Figure 2D are indicated. The mean value of each sample was calculated by Flowjo software.

**Figure 3-figure supplement 1.**
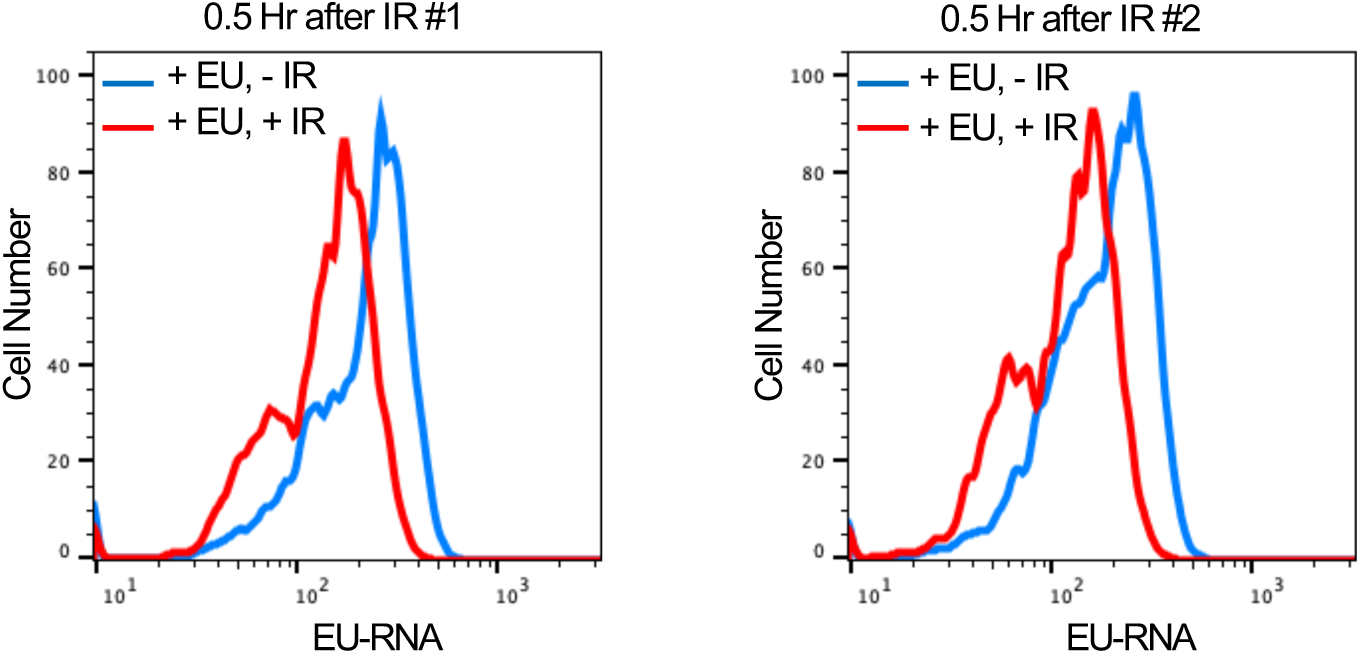
Nascent transcript levels before and after 30 minutes of IR. The samples from two independent experiments were used for the EU-seq and analyses in Fig. 3 and 4.

**Figure 3-figure supplement 2.**
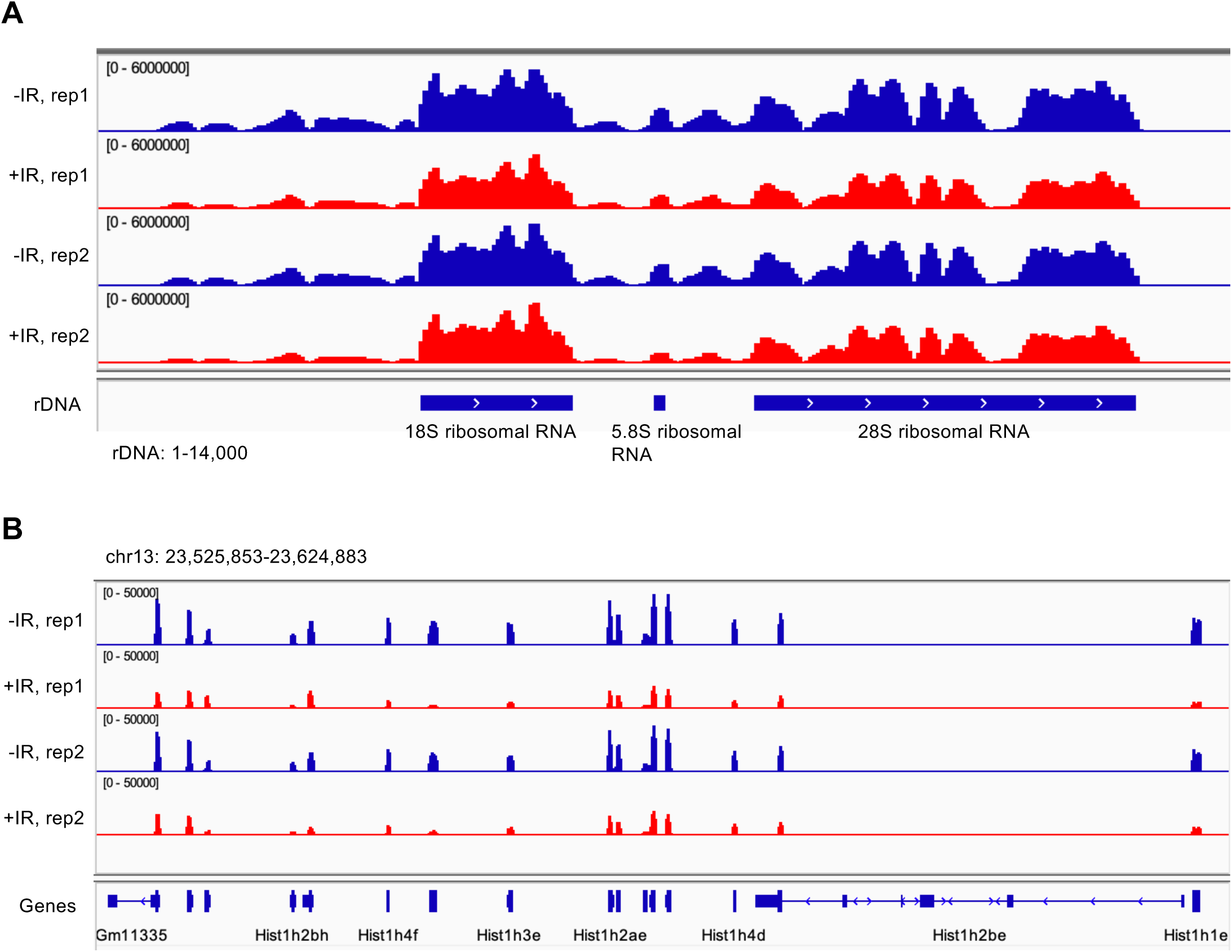
Screen shot from the UCSC browser of nascent transcripts. **A.** Nascent transcripts before and after IR over ribosomal DNA (rDNA). B. Nascent histone transcripts before and after IR over histone cluster 1.

**Figure 3-figure supplement 3.**
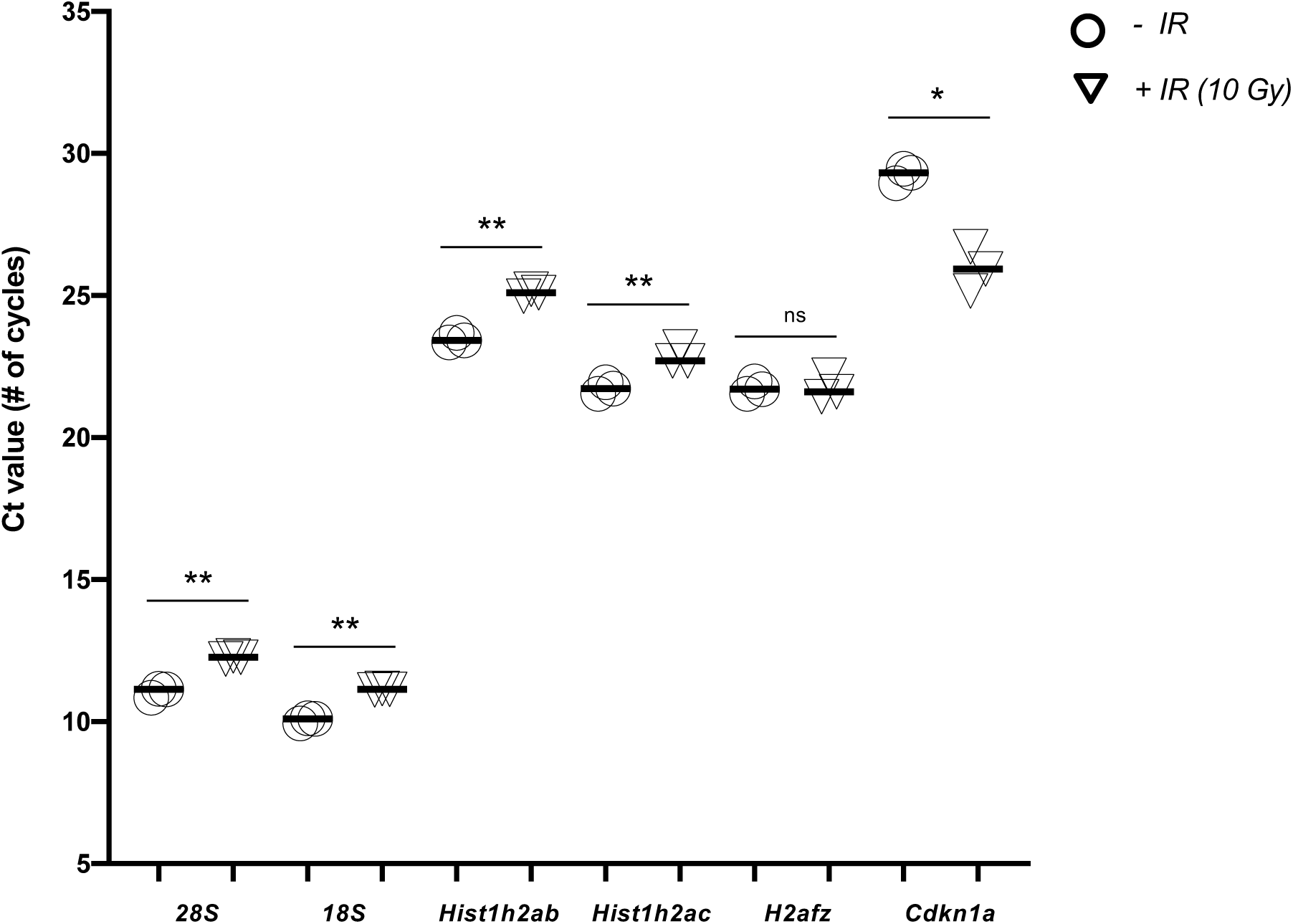
Validation of nascent transcript levels of DEGs from EU-RNA seq by real-time quantitative RT-PCR. Samples from three independent experiments were used for the analyses. Ct value is presented to show the absolute amounts of EU labeled RNA transcripts from the same number of cells before and after 30 minutes of IR (10 gray). Significant difference after IR compared to before IR are indicated by asterisks, where ** indicates p < 0.01, * indicates p < 0.05 and ns indicates non-significant by students T-test.

**Figure 5-figure supplement 1.**
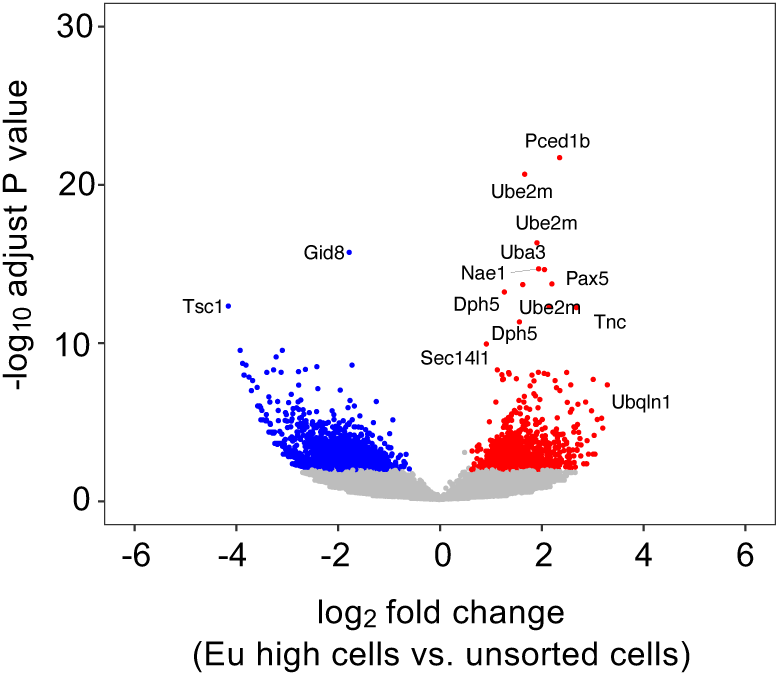
CRISPR-Cas9 screen identifies genes promoting transcriptional inhibition after IR. A volcano plot of guide RNA changes between Eu high cells and unsorted cells. Labeled genes are some of those that have P adjust <= 10^-6^.

**Figure 5-figure supplement 2.**
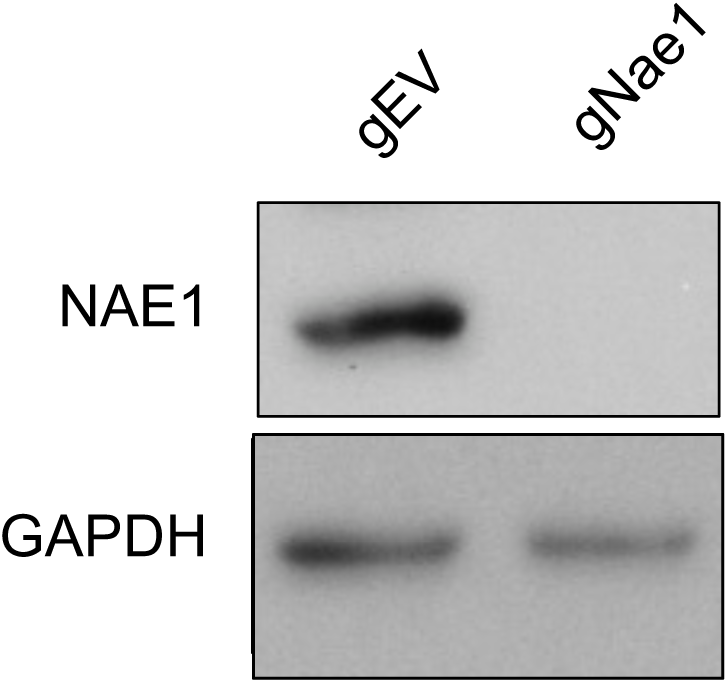
Confirmation of knockdown of Nae1. gEV is an empty vector. The samples were from the same experiments shown in Figure 5D.

**Figure 6-figure supplement 1.**
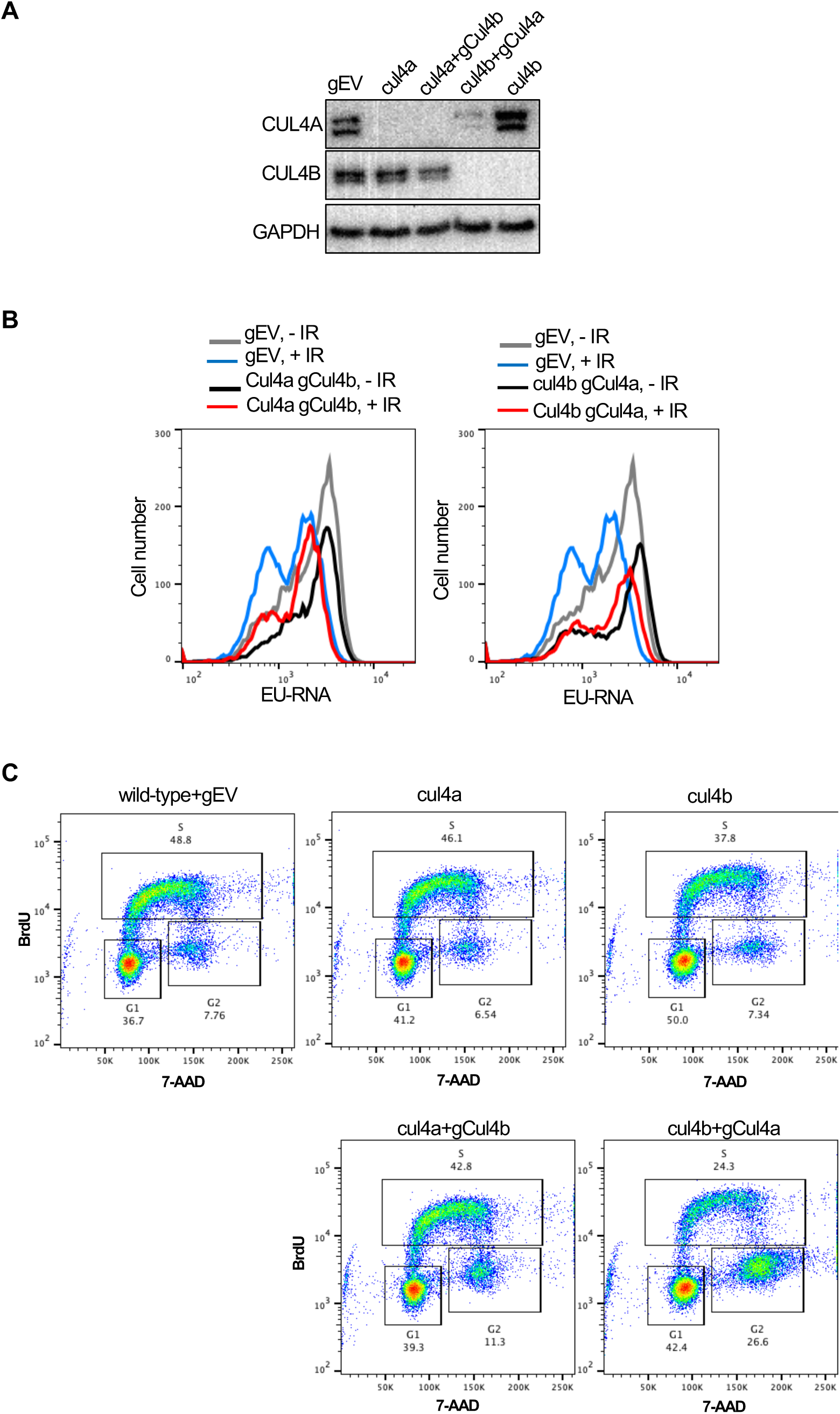
Analysis of CUL4A and CUL4B depletion. **A.** The western blot shows the CUL4A and CUL4B levels from the experiment shown in Fig. 6B. Additionally gRNAs were used to deplete CUL4A in cul4b cells and CUL4B in cul4a cells and their western blot analyses are also shown. **B.** The EU analysis of these double depleted cells is shown**. C.** Cell cycle analysis of the experiment shown in A and B and Fig. 6B.

**Figure 6-figure supplement 2.**
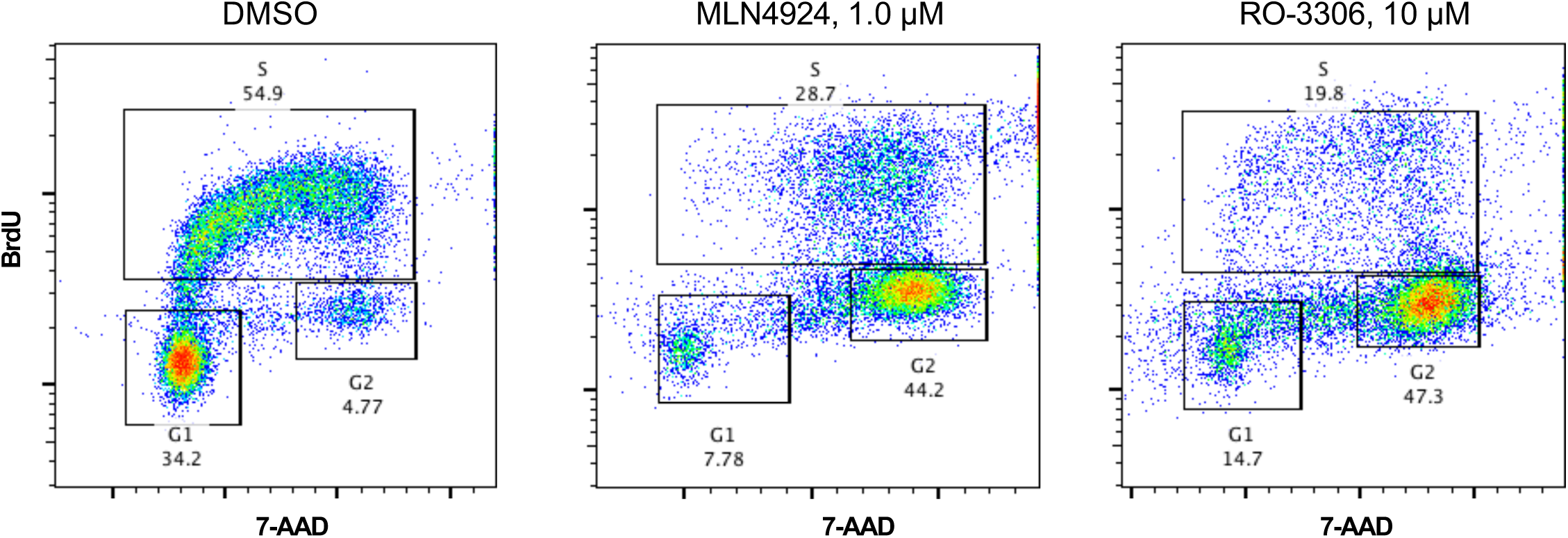
Cell cycle analysis of cells treated with MLN4924 and RO3306. MLN4924 or RO3306 treatment leads cell cycle arrest in G_2_ phase.

**Figure 7-figure supplement 1.**
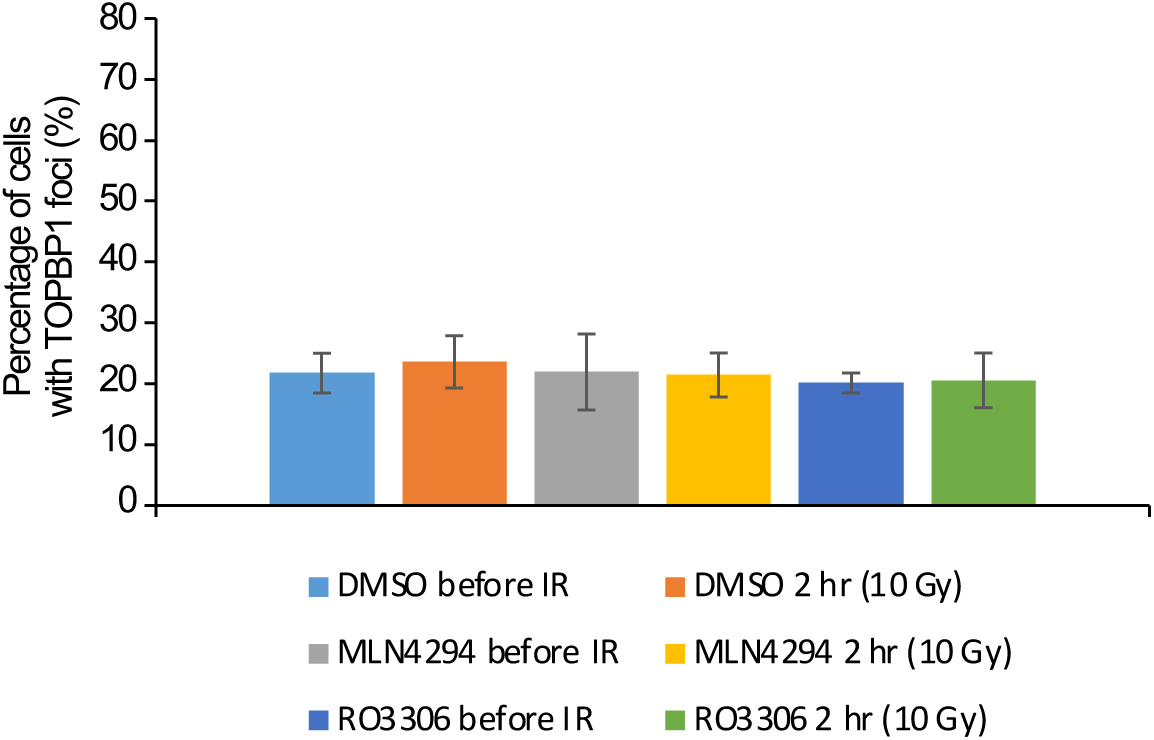
Quantification of cells with TOPBP1 expression before and after IR (10 Gy) in cells treated with MLN4924 and RO-3306. The samples were from the same experiments shown in Fig. 7F, 7G.

**Supplementary File 1.** Nascent RNA profiles of each gene using EU RNA-seq.

**Supplementary File 2.** Significantly enriched GO terms for up-regulated gene after irradiation.

**Supplementary File 3.** Significantly enriched GO terms for down-regulated genes after irradiation.

**Supplementary File 4.** Whole genome CRISPR-Cas9 screen detects the abundance of all gRNAs and target genes for EU high cells and unsorted cells.

**Figure 1-figure supplement 2 Source Data 1.** Original file for the Western blot analysis in Figure 1-Figure Supplement 1A (anti-HIRA and anti-GAPDH).

**Figure 1-figure supplement 2 Source Data 2.** PDF containing Figure 1-Figure Supplement 1A and original scans of the relevant Western blot analysis (anti-CUL4A, anti-CUL4B and anti-GAPDH) with highlighted bands and sample labels.

**Figure 1-figure supplement 2 Source Data 3**. Original file for the Western blot analysis in Figure 1-Figure Supplement 1B (anti-H3.3 and anti-GAPDH).

**Figure 1-figure supplement 2 Source Data 4.** PDF containing Figure 1-Figure Supplement 1B and original scans of the relevant Western blot analysis (anti-H3.3 and anti-GAPDH) with highlighted bands and sample labels.

**Figure 5-figure supplement 2 Source Data 1.** Original file for the Western blot analysis in Figure 5D (anti-Nae1 and anti-GAPDH).

**Figure 5-figure supplement 2 Source Data 2.** PDF of Western blot analysis in Figure 5D and original scans of the relevant Western blot analysis (anti-Nae1 and anti-GAPDH) with highlighted bands and sample labels.

**Figure 6 Source Data 1.** Original file for the Western blot analysis in Figure 6A (anti-CUL4A, anti-CUL4B and anti-GAPDH).

**Figure 6 Source Data 2.** PDF containing Figure 6A and original scans of the relevant Western blot analysis (anti-CUL4A, anti-CUL4B and anti-GAPDH) with highlighted bands and sample labels.

**Figure 6 Source Data 3.** Original file for the Western blot analysis in Figure 6C (anti-CUL4B, and anti-GAPDH).

**Figure 6 Source Data 4.** PDF containing Figure 6C and original scans of the relevant Western blot analysis (anti-CUL4B and anti-GAPDH) with highlighted bands and sample labels.

**Figure 6-figure supplement 1 Source Data 1.** Original file for the Western blot analysis in Figure 6-figure supplement 1A (anti-CUL4A, anti-CUL4B and anti-GAPDH).

**Figure 6-figure supplement 1 Source Data 2.** PDF containing Figure 6-figure supplement 1A and original scans of the relevant Western blot analysis (anti-CUL4A, anti-CUL4B and anti-GAPDH) with highlighted bands and sample labels.

## Methods and materials

### Cell culture and transfections

U2OS cells (ATCC, HTB-96) were cultured in McCoy’s 5A (Corning, 10050CV) medium supplemented with 10% fetal bovine serum (FBS) and 1% Penicillin-Streptomycin. Abelson virus-transformed pre-B cells were maintained in DMEM (Thermo Fisher, 11960-077) supplemented with 10% FBS, 1% Penicillin-Streptomycin, 1x nonessential amino acids, 1 mM sodium pyruvate, 2 mM L-glutamine, and 0.4% beta-mercaptoethanol. HEK-293T cells were maintained in DMEM (Corning, 10-013-CM) supplemented with 10% FBS and 1% Penicillin-Streptomycin. All the cells were grown at 37 °C under a humidified atmosphere with 5% CO2. SiRNA oligos against human H3F3A and H3F3B (SMARTPool) for RNAi in U2OS cells were purchased from Horizon Discovery (Dharmacon). 100 nM of H3F3A and H3F3B were mixed with Lipofectamine RNAiMAX transfection reagent (Thermo Scientific, 13778150) according to the manufacturer’s protocol to knockdown H3.3 for 48 hours. The siRNA control (ON-TARGETplus non-targeting) was also purchased from Horizon Discovery (Dharmacon) and used as negative control. shRNA lentiviral plasmids against HIRA (5’- TAGAGCATACCAAGATGCC-3’) and the control were described in a previous study (Huang et al., 2018). HEK-293T cells were transfected with a mixture of shRNA plasmids and the viral packaging and envelope vectors, pCMV-dR8.2 and pCMV-VSVG. The media containing shRNA virus particles were collected 48 to 72 hours after transfection and filtered through a 0.45 μm filter. Cells were incubated with the lentiviral supernatant containing 5 μg/ml polybrene (Sigma-Aldrich, S2667) for 24 hours, followed by 1 μg/mL Puromycin selection for another 48 hours. To inactivate Nae1, Cul4a and Cul4b in bulk cell populations, guide RNAs (gRNAs) against each gene were cloned into pKLV-U6gRNA-EF(BbsI)-PGKpuro2ABFP (Addgene, #62348) modified to express human CD2 or Thy1.1 as cell surface markers. The pKLV-gRNAs lentiviruses were prepared in 293T cells as described above. The Abl pre-B cells containing pCW-Cas9 (addgene, #50661), which can express Cas9 with doxycycline induction, were mixed with viral supernatant supplemented with 5 μg/ml polybrene and centrifuged at 1800 rpm for 1.5 hours at room temperature. After the spin-infection, the transduced cells were maintained in DMEM with 3 μg/ml doxycycline (Sigma-Aldrich, D9891) for 3 days before flow cytometric cell sorting based on hCD2 or Thy1.1 expression. To make stable cell lines depleted of Cul4a and Cul4b, serial dilution of the sorted cells into 96-well plate was used to isolate single cells. Western blot analysis was used to determine the knockdown efficiency of each target genes.

In EU flow cytometric and immunofluorescence analysis, final concentrations of 15 μM ATM inhibitor Ku55933 (Selleck Chemicals, S1092) and 5 μM of Actinomycin D (Sigma-Aldrich, A9415) were added to cell culture 1 hour prior to irradiation. 1 μM and 10 μM of neddylation inhibitor MLN4924/Pevonedistat (Active Biochem, A-1139) were used for long (16 hours) and short (1 to 3 hour) treatments, respectively. To arrest cells in G_1_ cell cycle phase, cells were incubated in media supplemented with 5 μM Palbociclib (Selleck Chemicals, S1116) or 5 μM Ribociclib (Selleck Chemicals, S7440) for 24 hours. For serum starvation, Abl pre-B cells were grown in complete medium containing 10% of FBS to desired density and collected and washed in medium with reduced concentration (0.1%) of FBS. The cells were maintained in the medium with reduced FBS for 72 hours to arrest in G_1_ phase. Representative data are shown for experiments repeated three of more times with consistent results.

U2OS cell lines were authenticated by STR profiling, and MCF10A and murine cell lines tested negative for mycoplasma contamination.

### Western blots

The following antibodies were used for western blot: CUL4A (Cell Signaling Technology, 2699S, 1:1000), CUL4B (Proteintech, 12916-1-AP, 1:1000), H3.3 (Millipore Sigma, 09-838, 1:1000), HIRA (Abcam, ab20655, 1:1000), NAE1 (Thermo Fisher, PA5-59836, 1:500), GAPDH (Sigma-Aldrich, G8795, 1:5000). Representative data are shown for experiments repeated three of more times with consistent results.

### Fluorescence microscopy

For immunofluorescence, Click-iT RNA Alexa Fluor Imaging Kit (Thermo Fisher, C10330) was used to label newly synthesized RNAs in the cells. Briefly, 50,000 U2OS cells grown on cover slips in 24-well plate were irradiated with 10 Gray and allowed to recover for indicated times at 37°C with 5% CO2. 0.5 mM EU was added to the medium and incubated for 45 minutes for EU incorporation. Cells were then washed with PBS, fixed in 4% paraformaldehyde PBS for 15 minutes at room temperature, and permeabilized in cold 0.5% Triton X-100 PBS for 10 minutes. Cells were blocked in 3% BSA-PBS for 1 hour at room temperature and subsequently incubated overnight at 4 °C in primary antibody (anti-γH2AX (S139), EMD Millipore, 05-636). Coverslips were then washed 3x with PBST (0.05% Tween 20), incubated with secondary antibody diluted in 3% BSA PBS (Alexa Fluor 488 Goat anti-mouse IgG, BioLegend, 405319) in the dark for 1 hour at room temperature and washed 3x with PBST. Click-iT reaction cocktail was prepared according to the manufacturer’s protocol and immediately added to the cells to perform click reaction in the dark for 30 minutes at room temperature. After washes with Click-iT reaction rinse buffer (Component F) and PBS, cells were stained with Hoechst (1:2000) or DAPI (Sigma-Aldrich, D9542) in PBS for 10 minutes and mounted in Prolong Gold Antifade Mountant (Life Technologies, P-36930). Images were taken on Biotek Lionheart Automatic Microscope and EU intensity quantification was conducted using Biotek Gen5 software. For eGFP-TOPBP1 fluorescence microscopy, 50,000 cells grown on coverslips in 24-well plates were treated with doxycycline (1 μg/mL) for 12 hours to express eGFP-TOPBP1. MLN4924 (1 μM) and RO3306 (10 μM) were added to the cell culture, and the cells were incubated for another 16 hours followed by IR (10 Gy) and recovery for 2 hours. Cells were fixed in 4% Paraformaldehyde for 20 minutes followed by Hoechst (1:2000) staining for DNA and mounting in Prolong Gold Antifade Mountant. Images were taken and quantified on Biotek Lionheart Automatic Microscope. Representative data are shown for experiments repeated three of more times with consistent results.

### Flow cytometry and cell cycle analysis

Click-iT RNA Alexa Fluor Imaging Kit was adapted to label newly synthesized RNAs in the cells for flow cytometry. Abl pre-B cells grown in 24-well plate were irradiated with 10 Gray and allowed to recover for different times. 2 mM EU was added to the medium and incubated for 30 minutes for EU incorporation. Cells were then washed with PBS, fixed in 4% paraformaldehyde PBS for 15 minutes at room temperature, and permeabilized in cold 0.5% TritonX-100 PBS for 5 minutes. Click-iT reaction cocktail was prepared according to the manufacturer’s protocol and immediately added to the cells to perform click reaction in the dark for 30 minutes at room temperature. Cells were then washed with Click-iT reaction rinse buffer (Component F) and 3% BSA-PBS, respectively. For cell cycle analysis, BrdU (10 ug/mL) was added to the cells and incubated for 30 minutes to label new DNAs. Cells were washed with PBS, fixed in 4% paraformaldehyde PBS for 15 minutes at room temperature, and permeabilized in cold 0.5% Triton X-100 PBS for 5 minutes. Cells were then digested with DNase (BD Biosciences, 51-2358KC) for 1 hour at 37 °C. Subsequently, cells were incubated with Alexa Fluor 488 Mouse anti-BrdU (BD Biosciences, 51-9004981, 1:500) in 3% BSA-FBS for 1 hour at room temperature and washed 2x with PBS, followed by FxCycle Violet (Thermo Scientific, R37166) or 7-AAD (BD Pharmingen, 559925) staining for 10 minutes. Cells were resuspended in PBS and analyzed on BD LSRII Flow Cytometer or BD LSRFortessa Flow Cytometer. Flow cytometry results were further analyzed using FlowJo software. Representative data are shown for experiments repeated three of more times with consistent results.

### CRISPR-Cas9 screen

More than 140 million wild type Abl pre-B cells carrying inducible Cas9 transgene were transduced with a lentiviral gRNA library containing 90,230 gRNAs targeting over 18,000 mouse genes (Addgene, 67988) by spin-infection as described above. 3 days post infection, cells transduced with gRNAs were sorted on a BD FACSAria II Cell Sorter based on BFP expression. BFP positive cells were treated 3 µg/ml doxycycline for 7 days to induce gRNA expression and gene inactivation. Cells were irradiated with 10 Gray, allowed to recover for 30 minutes, processed as described above for EU labeling of newly synthesized RNAs and analyzed on BD FACSAria II Cell Sorter. Cells with high (top 10%) EU staining and unsorted cells were collected, and genomic DNA of the cells were isolated for library preparation using nested-PCR. The library was sequenced on an Illumina HiSeq 2500 platform. Raw fastq files were demultiplexed by the Genomics and Epigenomics Core Facility of the Weill Cornell Medicine Core Laboratories Center. The gRNA sequence region was then retrieved from the sequencing data using Seqkit (Shen et al., 2016) and mapped to the gRNA sequence library (Koike-Yusa et al., 2014, Tzelepis et al., 2016). The number of reads of each library sequence was counted and then normalized as follows (Shalem et al., 2014). Normalized reads per gRNA = reads pers gRNA total reads for all sgRNAs in sample×10^6^+1. Hereby, the generated normalized reads from each guide RNA were used and compared between the EU high cell and unsorted cell. P values were measured by Poisson test to compare guide RNAs between EU high cell and unsorted cell. FDR was used for adjusting P value. CRISPR score = log2 (final sgRNA abundance/initial sgRNA abundance). The EU high genes were defined as these genes that have at least one guide RNA with P adjust value <=0.01 & FC >=1.5. Gene ontology (GO) analysis was performed by the R package cluster Profiler v3.18.1.

### Isolation and deep sequencing of EU labeled nascent transcripts

Click-iT Nascent RNA Capture Kit (Thermo Fisher, C10365) was used to label and capture the nascent transcripts. In brief, the same number of Abl pre-B cells was plated in two T-25 Polystyrene flasks, one for irradiation (10 Gray) and the other for no IR control. After irradiation, both flasks of cells were recovered for 30 minutes and incubated in medium with 2 mM EU for 30 minutes to allow for the incorporation of EU into the nascent transcripts. Total RNAs were harvested using TRIzol reagent (Thermo Fischer, 15596018) following the manufacturer’s protocol. The click-iT reaction was performed as per manufacturer’s protocol in 50 µL total volume for 30 minutes in the dark. Subsequently, 1 µL ultrapure glycogen, 50 µL 7.5 M ammonium acetate, and 700 µL of chilled 100% EtOH were added to the reaction. The mixture was incubated at −80 °C for 16 hours. RNA pellet was spun down at 13000 ×g for 20 minutes at 4 °C, washed 2x with chilled 75% EtOH and resuspended in nuclease free water. EU-RNAs were pulled down with Dynabeads MyOne Streptavidin T1 magnetic beads and extracted with TRIzol reagents. The same amount of ERCC spike-ins (Thermo Fisher, 4456740) were added to the purified EU-RNAs, followed by cDNA library generation using NEBNext Ultr II Directional RNA Library Prep Kit for Illumina (NEB, E7760) according to the manufacturer’s protocol and deep sequencing on Illumina HiSeq 2500 platform.

### Real-time quantitative RT-PCR of EU labeled nascent transcripts

EU labeled RNA was prepared from the same number of cells as described above. All the isolated EU-RNAs were used for cDNA synthesis using Superscript III (Thermo Fischer, 18080044) reverse transcription with random hexamer as primers following the manufacturer’s protocol. The same proportion of cDNA products of each sample was used as template for the quantitative RT-PCR reaction with Light Cycler 480 SYBR Green I Master Mix (Roche, 04707516001). The Ct value was used to represent the absolute amount of EU-RNAs in each sample, in which a smaller Ct value indicates higher nascent transcript levels of an individual gene, given the same initial number of cells and the same proportion of EU-RNAs were used for the analyses. Primer sequences for the analyses are as follows:

ms28S-fwd, 5’-TGGGTTTTAAGCAGGAGGTG-; ms28S-rev, 5’- GTGAATTCTGCTTCACAATG-3’(Watada et al., 2020); ms18S-fwd, 5’- CTTAGAGGGACAAGTGGCG-3’; ms18S-rev, 5’-ACGCTGAGCCAGTCAGTGTA-3’ (StephensStephens and Morrison, 2011); msHist1h2ab-fwd, 5’-GCCTGCAGTTCCCCGTA-3’; msHisth2ab-rev, 5’- ATCTCGGCCGTCAGGTACTC-3’; msHist1h2ac-fwd, 5’- GGCTGCTCCGCAAGGGT-3’; msHist1h2ac-rev, 5’-CTTGTTGAGCTCCTCGTCGTT-3’; msH2afz-fwd, 5’- ACTCCGGAAAGGCCAAGACA-3’; msH2afz-rev, 5’- GTTGTCCTAGATTTCAGGTG-3’ (Nishida et al., 2005); msCdkn1a-fwd, 5’- GTGGCCTTGTCGCTGTCT-3’; msCdkn1a-rev, 5’-TTTTCTCTTGCAGAAGACCAATC-3’ (Béguelin et al., 2017).

### Analysis of nascent transcripts

The mouse genome version GRCm38.p6 release M23 and the associated GENCODE version of mouse reference gene set were downloaded from the GENCODE website (https://www.gencodegenes.org/mouse/release_M23.html). We trimmed adapter sequences and low-quality sequences in RNA-seq data using the Trim Galore v0.6.6 (Martin, 2011) with default parameters. To avoid rRNA homologous sequences (i.e., in the intron regions of Zc3h7a or Cdk8) prior to subsequent genomic and other RNA analysis, we first mapped the reads to mm10 rDNA sequences by TopHat v2.1.1 (Kim et al., 2013). The unmapped reads were then further mapped to the mouse genome version mm10 and ERCC spike-in version ERCC92 using TopHat v2.1.1 (Kim et al., 2013). Successfully mapping reads were sorted by SAMtools v1. 5. Afterward, read counts in several types of genomic feature, i.e., protein-coding genes, rDNA and ERCCs (ERCC92.gtf), were quantified by Htseq-count v0.11.2 (AndersPyl and Huber, 2015) using the union gene region option. The read number per gene was normalized based on total ERCC read numbers in each sample.

To visualize read coverage across the genome, DeepTools v3.5.0 (Ramírez et al., 2014) was used to convert BAM files into bigwig files using scale factors calculated by total ERCC read number in each sample. Next, DeepTools was used to plot average read depth per sample across interested groups of genomic regions (i.e., repressed protein coding genes from 3 kb upstream to 3kb downstream of gene bodies). Screenshots of reads density at individual regions were generated by IGV 2.8.13 (ThorvaldsdóttirRobinson and Mesirov, 2013).

We then used one tail Poisson test to evaluate difference in gene expression level based on the read counts normalized by total ERCC read counts. We defined differentially expressed RNAs as those with a fold change greater than 1.5 and an FDR value smaller than 0.05. To detect highly expressed genes, we ranked genes by RPKM in the control cells, whereas RPKM was calculated using ERCC-normalized read counts further normalized by gene length. Gene ontology (GO) analysis was performed by the R package clusterProfiler v3.18.1 (Wu et al., 2021). Heatmap were generated by pheatmap.

## Data availability

All raw sequencing data from the nascent EU RNA-seq and CRISPR screen experiments have been deposited in the NCBI project database under accession PRJNA895065. The genome-wide gRNA library CRISPR-Case9 screen datasets comprise Abl pre-B cells of both unsorted (SRX18076832) and sorted the 10% of the cells with the most nascent RNA (high EU) 30 minutes after IR (SRX18076831). The raw FASTQ files for nascent EU RNA-seq include pre-B cells without irradiation (SRX18050529 and SRX18050531) and with 30 minutes after irradiation (SRX18050530 and SRX18050532). Additionally, both FASTQ files and processed data for nascent EU RNA-seq are accessible at GSE217123. The source codes employed in the data analysis and figure generation have been uploaded to GitHub at the following repository: https://github.com/gucascau/NascentDiff.git.

The following data sets were generated:

Chen et al (2024) NCBI BioProject ID PRJNA895065. Transcriptional inhibition after irradiation occurs preferentially at highly expressed genes in a manner dependent on cell cycle progression https://www.ncbi.nlm.nih.gov/bioproject/PRJNA895065

Chen et al (2024) NCBI Gene Expression Omnibus, GSE217123. Transcriptional inhibition after irradiation occurs preferentially at highly expressed genes in a manner dependent on cell cycle progression https://www.ncbi.nlm.nih.gov/geo/query/acc.cgi?acc=GSE217123

## Acknowledgements

The authors thank Yinan Wang for performing the bioinformatics for the high throughput screen. We thank the Weill Cornell Flow Cytometry Core for flow cytometry. We thank the Weill Cornell Epigenomics Core for performing the sequencing for the high throughput screen and the Transcriptional Regulation & Expression Facility at Cornell University, Ithaca for providing advice and performing the sequencing of the nascent transcripts. The stable doxycycline-inducible eGFP-TOPBP1 U2OS cell lines were a kind gift from Dr. Helmut Pospiech (Fritz Lipmann Institute, Germany). We are grateful to Pengbo Zhou for advice on Cullin 4A and B. We also thank Barry Sleckman, Bo-Ruei Chen and Faith Fowler for advice throughout the project. JKT is supported by NIH R35 GM139816 and RO1 CA95641. K.C is supported by NIH R01GM138407, R01GM125632, R01HL148338, and R01HL133254.

## Notes

### Competing Interest Statement

The authors have declared no competing interest.

### Summary of Updates

The version has been revised in response to reviews from eLife

## References

Adam, S., Polo, S. E. & Almouzni, G. 2013. Transcription recovery after DNA damage requires chromatin priming by the H3.3 histone chaperone HIRA. Cell, 155, 94–106.

Anders, S., Pyl, P. T. & Huber, W. 2015. HTSeq—a Python framework to work with high-throughput sequencing data. bioinformatics, 31, 166–169.

Arnould, C., Rocher, V., Saur, F., Bader, A. S., Muzzopappa, F., Collins, S., Lesage, E., le Bozec, B., Puget, N., Clouaire, T., Mangeat, T., Mourad, R., Ahituv, N., Noordermeer, D., Erdel, F., Bushell, M., Marnef, A. & Legube, G. 2023. Author Correction: Chromatin compartmentalization regulates the response to DNA damage. Nature, 624, E1.

Artuso, M., Esteve, A., Bresil, H., Vuillaume, M. & Hall, J. 1995. The role of the Ataxia telangiectasia gene in the p53, WAF1/CIP1(p21)- and GADD45-mediated response to DNA damage produced by ionising radiation. Oncogene, 11, 1427–35.

Béguelin, W., Rivas, M. A., Calvo Fernández, M. T., Teater, M., Purwada, A., Redmond, D., Shen, H., Challman, M. F., Elemento, O., Singh, A. & Melnick, A. M. 2017. EZH2 enables germinal centre formation through epigenetic silencing of CDKN1A and an Rb-E2F1 feedback loop. Nat Commun, 8, 877.

Blackford, A. N. & Jackson, S. P. 2017. Atm, Atr, and DNA-PK: The Trinity at the Heart of the DNA Damage Response. Mol Cell, 66, 801–817.

Bouvier, D., Ferrand, J., Chevallier, O., Paulsen, M. T., Ljungman, M. & Polo, S. E. 2021. Dissecting regulatory pathways for transcription recovery following DNA damage reveals a non-canonical function of the histone chaperone HIRA. Nat Commun, 12, 3835.

Bredemeyer, A. L., Sharma, G. G., Huang, C. Y., Helmink, B. A., Walker, L. M., Khor, K. C., Nuskey, B., Sullivan, K. E., Pandita, T. K., Bassing, C. H. & Sleckman, B. P. 2006. ATM stabilizes DNA double-strand-break complexes during V(D)J recombination. Nature, 442, 466–70.

Brown, J. S., Lukashchuk, N., Sczaniecka-Clift, M., Britton, S., le Sage, C., Calsou, P., Beli, P., Galanty, Y. & Jackson, S. P. 2015. Neddylation promotes ubiquitylation and release of Ku from DNA-damage sites. Cell Rep, 11, 704–14.

Caron, P., Pankotai, T., Wiegant, W. W., Tollenaere, M. A. X., Furst, A., Bonhomme, C., Helfricht, A., de Groot, A., Pastink, A., Vertegaal, A. C. O., Luijsterburg, M. S., Soutoglou, E. & van Attikum, H. 2019. WWP2 ubiquitylates RNA polymerase II for DNA-PK-dependent transcription arrest and repair at DNA breaks. Genes Dev, 33, 684–704.

Chen, B. R., Wang, Y., Tubbs, A., Zong, D., Fowler, F. C., Zolnerowich, N., Wu, W., Bennett, A., Chen, C. C., Feng, W., Nussenzweig, A., Tyler, J. K. & Sleckman, B. P. 2021. LIN37-DREAM prevents DNA end resection and homologous recombination at DNA double-strand breaks in quiescent cells. Elife, 10.

Chen, K., Hu, Z., Xia, Z., Zhao, D., Li, W. & Tyler, J. K. 2015. The Overlooked Fact: Fundamental Need for Spike-In Control for Virtually All Genome-Wide Analyses. Mol Cell Biol, 36, 662–7.

Hannah, J. & Zhou, P. 2015. Distinct and overlapping functions of the cullin E3 ligase scaffolding proteins CUL4A and CUL4B. Gene, 573, 33–45.

Huang, T. H., Fowler, F., Chen, C. C., Shen, Z. J., Sleckman, B. & Tyler, J. K. 2018. The Histone Chaperones ASF1 and CAF-1 Promote MMS22L-TONSL-Mediated Rad51 Loading onto ssDNA during Homologous Recombination in Human Cells. Mol Cell, 69, 879–892 e5.

Iannelli, F., Galbiati, A., Capozzo, I., Nguyen, Q., Magnuson, B., Michelini, F., D’alessandro, G., Cabrini, M., Roncador, M., Francia, S., Crosetto, N., Ljungman, M., Carninci, P. & D’adda di Fagagna, F. 2017. A damaged genome’s transcriptional landscape through multilayered expression profiling around in situ-mapped DNA double-strand breaks. Nat Commun, 8, 15656.

Jackson, S. P. & Bartek, J. 2009. The DNA-damage response in human biology and disease. Nature, 461, 1071–8.

Jao, C. Y. & Salic, A. 2008. Exploring RNA transcription and turnover in vivo by using click chemistry. Proc Natl Acad Sci U S A, 105, 15779–84.

Kakarougkas, A., Ismail, A., Chambers, A. L., Riballo, E., Herbert, A. D., Kunzel, J., Lobrich, M., Jeggo, P. A. & Downs, J. A. 2014. Requirement for PBAF in transcriptional repression and repair at DNA breaks in actively transcribed regions of chromatin. Mol Cell, 55, 723–32.

Kim, D., Pertea, G., Trapnell, C., Pimentel, H., Kelley, R. & Salzberg, S. L. 2013. TopHat2: accurate alignment of transcriptomes in the presence of insertions, deletions and gene fusions. Genome biology, 14, 1–13.

Kim, J. H., Jenrow, K. A. & Brown, S. L. 2014. Mechanisms of radiation-induced normal tissue toxicity and implications for future clinical trials. Radiat Oncol J, 32, 103–15.

Koike-Yusa, H., Li, Y., Tan, E. P., Velasco-Herrera, M. E. C. & Yusa, K. 2014. Genome-wide recessive genetic screening in mammalian cells with a lentiviral CRISPR-guide RNA library. Nat Biotechnol, 32, 267–73.

Kruhlak, M., Crouch, E. E., Orlov, M., Montano, C., Gorski, S. A., Nussenzweig, A., Misteli, T., Phair, R. D. & Casellas, R. 2007. The ATM repair pathway inhibits RNA polymerase I transcription in response to chromosome breaks. Nature, 447, 730–4.

Larsen, D. H., Hari, F., Clapperton, J. A., Gwerder, M., Gutsche, K., Altmeyer, M., Jungmichel, S., Toledo, L. I., Fink, D., Rask, M. B., Grøfte, M., Lukas, C., Nielsen, M. L., Smerdon, S. J., Lukas, J. & Stucki, M. 2014. The NBS1-Treacle complex controls ribosomal RNA transcription in response to DNA damage. Nat Cell Biol, 16, 792–803.

Lieberman, H. B., Panigrahi, S. K., Hopkins, K. M., Wang, L. & Broustas, C. G. 2017. p53 and Rad9, the DNA Damage Response, and Regulation of Transcription Networks. Radiat Res, 187, 424–432.

Ma, T., van Tine, B. A., Wei, Y., Garrett, M. D., Nelson, D., Adams, P. D., Wang, J., Qin, J., Chow, L. T. & Harper, J. W. 2000. Cell cycle-regulated phosphorylation of p220(NPAT) by cyclin E/Cdk2 in Cajal bodies promotes histone gene transcription. Genes Dev, 14, 2298–313.

Martin, M. 2011. Cutadapt removes adapter sequences from high-throughput sequencing reads. EMBnet. journal, 17, 10–12.

Meisenberg, C., Pinder, S. I., Hopkins, S. R., Wooller, S. K., Benstead-Hume, G., Pearl, F. M. G., Jeggo, P. A. & Downs, J. A. 2019. Repression of Transcription at DNA Breaks Requires Cohesin throughout Interphase and Prevents Genome Instability. Mol Cell, 73, 212–223 e7.

Mooser, C., Symeonidou, I. E., Leimbacher, P. A., Ribeiro, A., Shorrocks, A. K., Jungmichel, S., Larsen, S. C., Knechtle, K., Jasrotia, A., Zurbriggen, D., Jeanrenaud, A., Leikauf, C., Fink, D., Nielsen, M. L., Blackford, A. N. & Stucki, M. 2020. Treacle controls the nucleolar response to rDNA breaks via TOPBP1 recruitment and ATR activation. Nat Commun, 11, 123.

Moss, T. & Stefanovsky, V. Y. 1995. Promotion and regulation of ribosomal transcription in eukaryotes by RNA polymerase I. Prog Nucleic Acid Res Mol Biol, 50, 25–66.

Nishida, H., Suzuki, T., Ookawa, H., Tomaru, Y. & Hayashizaki, Y. 2005. Comparative analysis of expression of histone H2a genes in mouse. BMC Genomics, 6, 108.

O’mahony, D. J., Xie, W. Q., Smith, S. D., Singer, H. A. & Rothblum, L. I. 1992. Differential phosphorylation and localization of the transcription factor UBF in vivo in response to serum deprivation. In vitro dephosphorylation of UBF reduces its transactivation properties. J Biol Chem, 267, 35–8.

Pan, Z. Q., Kentsis, A., Dias, D. C., Yamoah, K. & Wu, K. 2004. Nedd8 on cullin: building an expressway to protein destruction. Oncogene, 23, 1985–97.

Pankotai, T., Bonhomme, C., Chen, D. & Soutoglou, E. 2012. DNAPKcs-dependent arrest of RNA polymerase II transcription in the presence of DNA breaks. Nat Struct Mol Biol, 19, 276–82.

Pankotai, T. & Soutoglou, E. 2013. Double strand breaks: hurdles for RNA polymerase II transcription? Transcription, 4, 34–8.

Porter, J. R., Fisher, B. E., Baranello, L., Liu, J. C., Kambach, D. M., Nie, Z., Koh, W. S., Luo, J., Stommel, J. M., Levens, D. & Batchelor, E. 2017. Global Inhibition with Specific Activation: How p53 and MYC Redistribute the Transcriptome in the DNA Double-Strand Break Response. Mol Cell, 67, 1013–1025 e9.

Purman, C. E., Collins, P. L., Porter, S. I., Saini, A., Gupta, H., Sleckman, B. P. & Oltz, E. M. 2019. Regional Gene Repression by DNA Double-Strand Breaks in G1 Phase Cells. Mol Cell Biol, 39.

Rabut, G. & Peter, M. 2008. Function and regulation of protein neddylation. ‘Protein modifications: beyond the usual suspects’ review series. EMBO Rep, 9, 969–76.

Ramírez, F., Dündar, F., Diehl, S., Grüning, B. A. & Manke, T. 2014. deepTools: a flexible platform for exploring deep-sequencing data. Nucleic acids research, 42, W187–W191.

Schaue, D., Kachikwu, E. L. & Mcbride, W. H. 2012. Cytokines in radiobiological responses: a review. Radiat Res, 178, 505–23.

Shalem, O., Sanjana, N. E., Hartenian, E., Shi, X., Scott, D. A., Mikkelson, T., Heckl, D., Ebert, B. L., Root, D. E., Doench, J. G. & Zhang, F. 2014. Genome-scale CRISPR-Cas9 knockout screening in human cells. Science, 343, 84–87.

Shanbhag, N. M., Rafalska-Metcalf, I. U., Balane-Bolivar, C., Janicki, S. M. & Greenberg, R. A. 2010. ATM-dependent chromatin changes silence transcription in cis to DNA double-strand breaks. Cell, 141, 970–81.

Shen, W., Le, S., Li, Y. & Hu, F. 2016. SeqKit: A Cross-Platform and Ultrafast Toolkit for FASTA/Q File Manipulation. PLoS One, 11, e0163962.

Sittman, D. B., Graves, R. A. & Marzluff, W. F. 1983. Histone mRNA concentrations are regulated at the level of transcription and mRNA degradation. Proc Natl Acad Sci U S A, 80, 1849–53.

Sokka, M., Rilla, K., Miinalainen, I., Pospiech, H. & Syväoja, J. E. 2015. High levels of TopBP1 induce ATR-dependent shut-down of rRNA transcription and nucleolar segregation. Nucleic Acids Res, 43, 4975–89.

Stephens, A. S., Stephens, S. R. & Morrison, N. A. 2011. Internal control genes for quantitative RT-PCR expression analysis in mouse osteoblasts, osteoclasts and macrophages. BMC Res Notes, 4, 410.

Su, C., Gao, G., Schneider, S., Helt, C., Weiss, C., O’reilly, M. A., Bohmann, D. & Zhao, J. 2004. DNA damage induces downregulation of histone gene expression through the G1 checkpoint pathway. EMBO J, 23, 1133–43.

Thorvaldsdóttir, H., Robinson, J. T. & Mesirov, J. P. 2013. Integrative Genomics Viewer (IGV): high-performance genomics data visualization and exploration. Briefings in bioinformatics, 14, 178–192.

Tzelepis, K., Koike-Yusa, H., de Braekeleer, E., Li, Y., Metzakopian, E., Dovey, O. M., Mupo, A., Grinkevich, V., Li, M., Mazan, M., Gozdecka, M., Ohnishi, S., Cooper, J., Patel, M., Mckerrell, T., Chen, B., Domingues, A. F., Gallipoli, P., Teichmann, S., Ponstingl, H., Mcdermott, U., Saez-Rodriguez, J., Huntly, B. J. P., Iorio, F., Pina, C., Vassiliou, G. S. & Yusa, K. 2016. A CRISPR Dropout Screen Identifies Genetic Vulnerabilities and Therapeutic Targets in Acute Myeloid Leukemia. Cell Rep, 17, 1193–1205.

Van Sluis, M. & Mcstay, B. 2017. Nucleolar reorganization in response to rDNA damage. Curr Opin Cell Biol, 46, 81–86.

Vassilev, L. T., Tovar, C., Chen, S., Knezevic, D., Zhao, X., Sun, H., Heimbrook, D. C. & Chen, L. 2006. Selective small-molecule inhibitor reveals critical mitotic functions of human CDK1. Proc Natl Acad Sci U S A, 103, 10660–5.

Venkata Narayanan, I., Paulsen, M. T., Bedi, K., Berg, N., Ljungman, E. A., Francia, S., Veloso, A., Magnuson, B., DI Fagagna, F. D., Wilson, T. E. & Ljungman, M. 2017. Transcriptional and post-transcriptional regulation of the ionizing radiation response by ATM and p53. Sci Rep, 7, 43598.

Watada, E., Li, S., Hori, Y., Fujiki, K., Shirahige, K., Inada, T. & Kobayashi, T. 2020. Age-Dependent Ribosomal DNA Variations in Mice. Mol Cell Biol, 40.

Wu, T., Hu, E., Xu, S., Chen, M., Guo, P., Dai, Z., Feng, T., Zhou, L., Tang, W. & Zhan, L. 2021. clusterProfiler 4.0: A universal enrichment tool for interpreting omics data. The Innovation, 2, 100141.

Yu, Z., Xu, C., Song, B., Zhang, S., Chen, C., Li, C. & Zhang, S. 2023. Tissue fibrosis induced by radiotherapy: current understanding of the molecular mechanisms, diagnosis and therapeutic advances. J Transl Med, 21, 708.

Zhao, J., Dynlacht, B., Imai, T., Hori, T. & Harlow, E. 1998. Expression of Npat, a novel substrate of cyclin E-Cdk2, promotes S-phase entry. Genes Dev, 12, 456–61.

Zhao, J., Kennedy, B. K., Lawrence, B. D., Barbie, D. A., Matera, A. G., Fletcher, J. A. & Harlow, E. 2000. NPAT links cyclin E-Cdk2 to the regulation of replication-dependent histone gene transcription. Genes Dev, 14, 2283–97.

