## Supplementary material for "Transcriptional inhibition after irradiation occurs preferentially at highly expressed genes in a manner dependent on cell cycle progression": Source data

### Slide 1
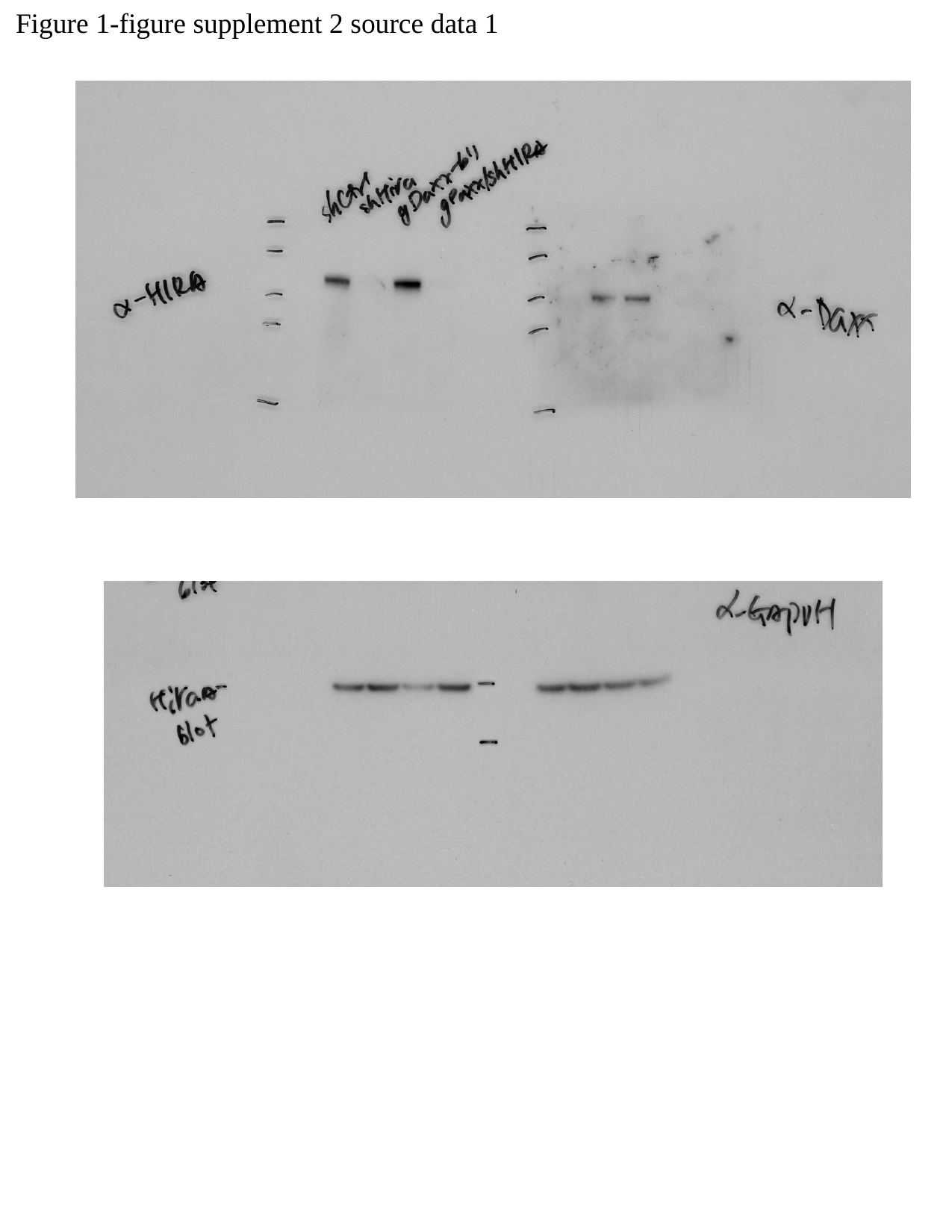

Figure 1-figure supplement 2 source data 1

### Slide 2
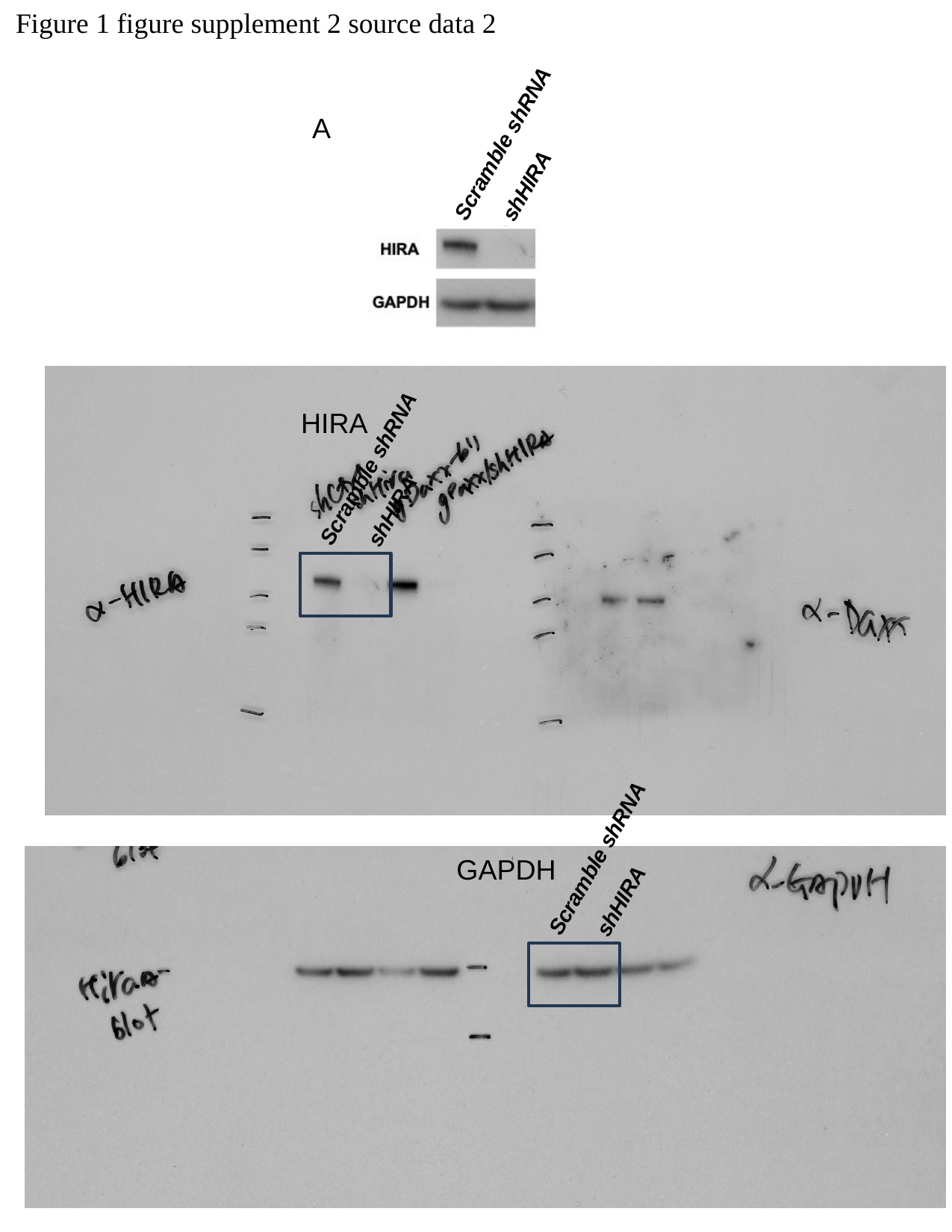

Figure 1 figure supplement 2 source data 2
A
Scramble shRNA
shHIRA
HIRA
Scramble shRNA
shHIRA
Scramble shRNA
GAPDH
shHIRA

### Slide 3
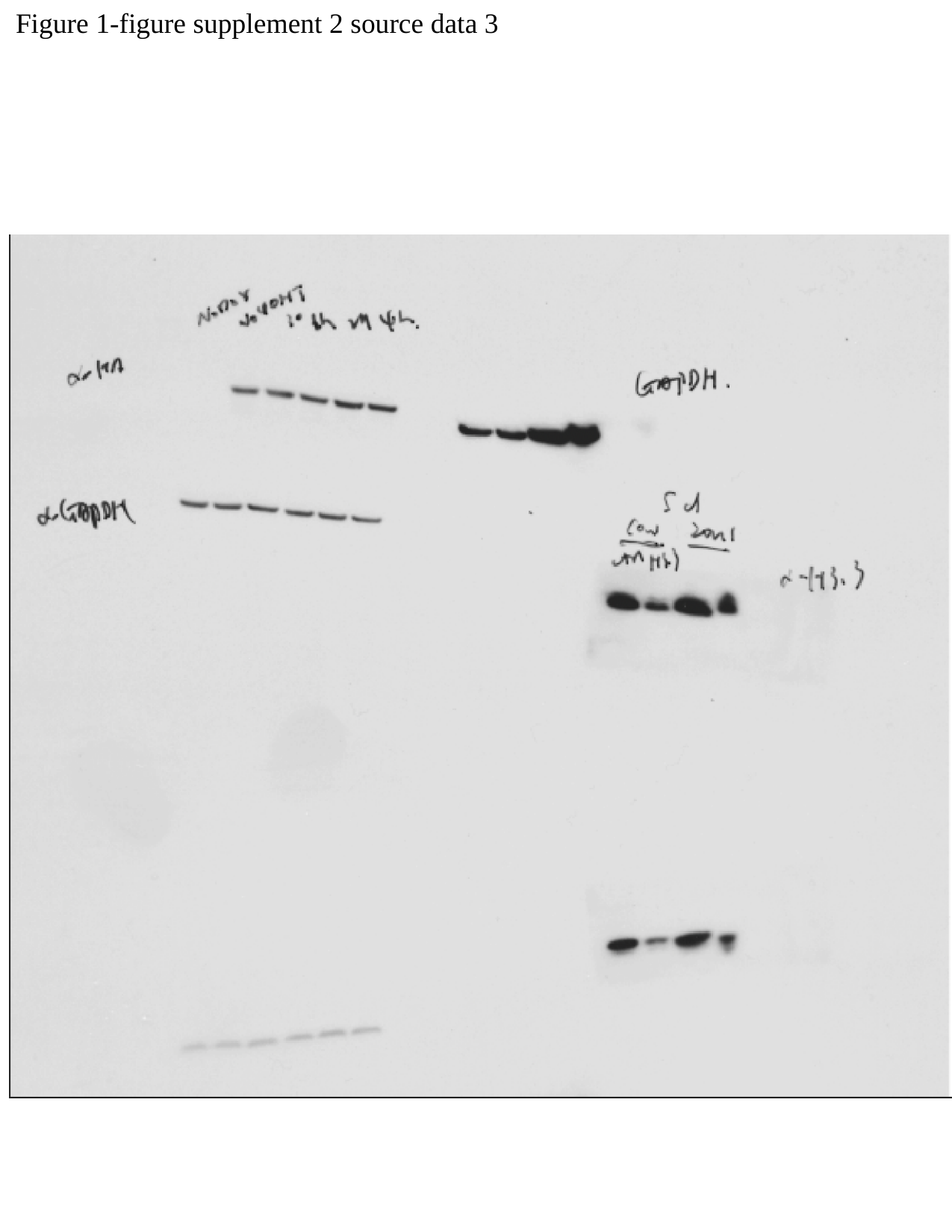

Figure 1-figure supplement 2 source data 3

### Slide 4
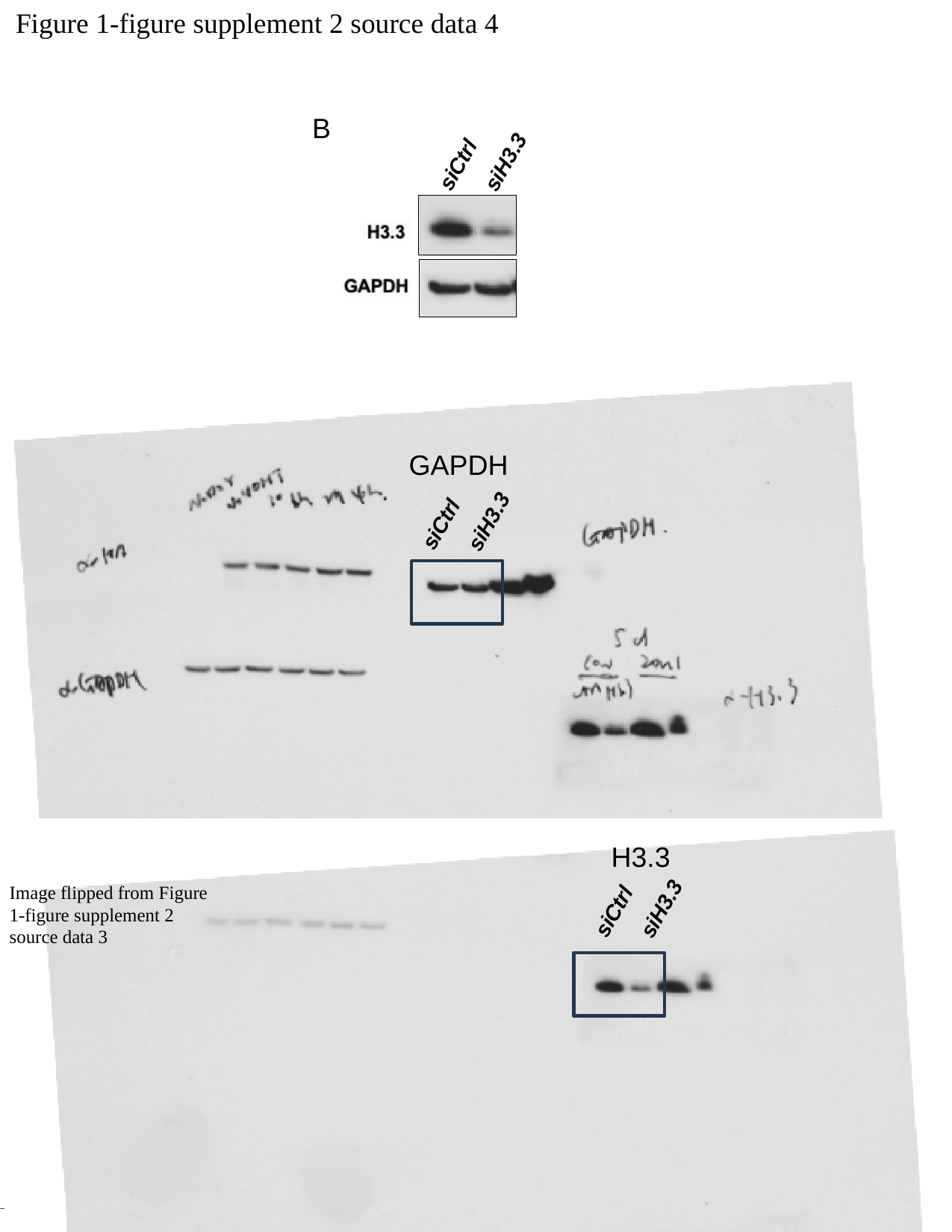

Figure 1-figure supplement 2 source data 4
B
siCtrl
siH3.3
GAPDH
siCtrl
siH3.3
H3.3
Image flipped from Figure 1-figure supplement 2 source data 3
siCtrl
siH3.3

### Slide 5
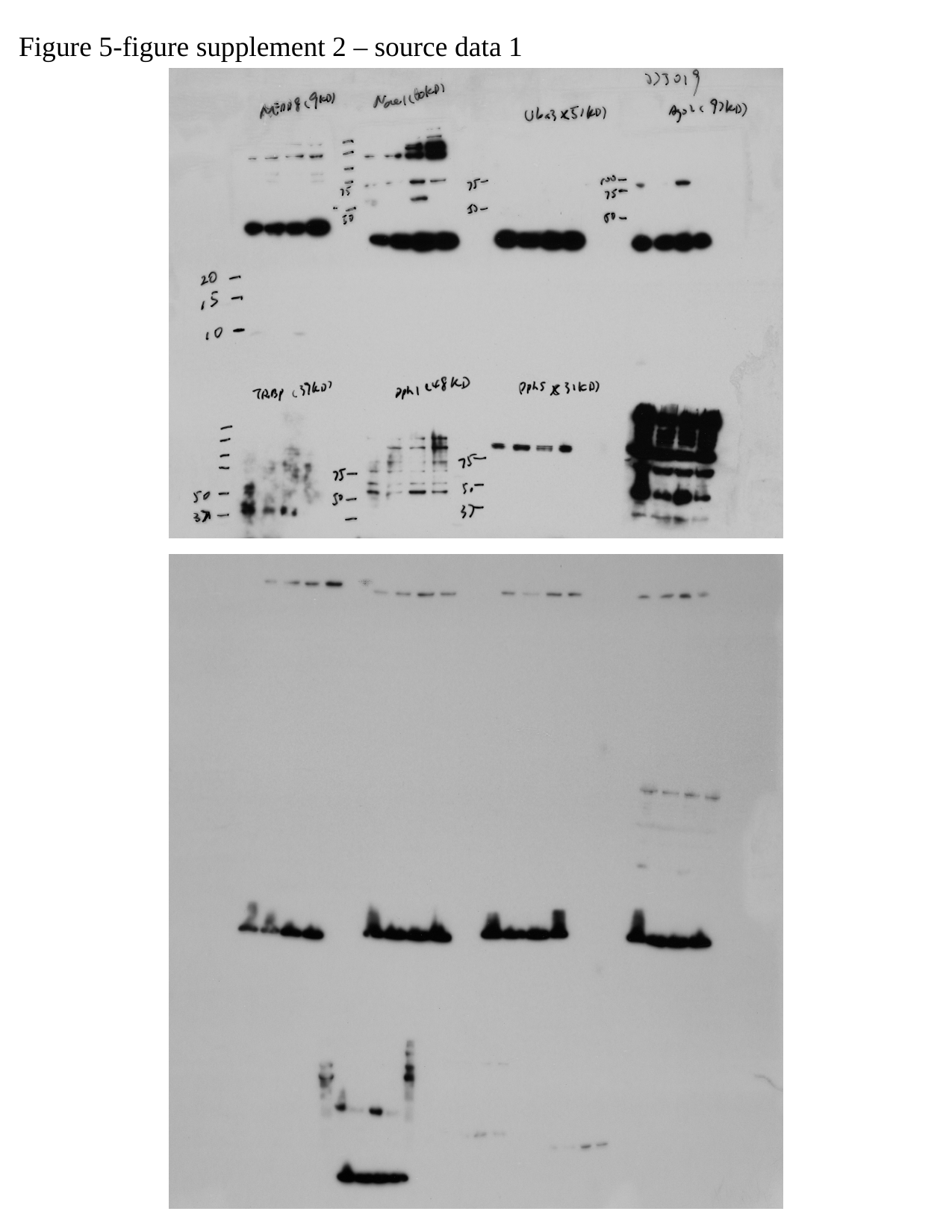

Figure 5-figure supplement 2 – source data 1

### Slide 6
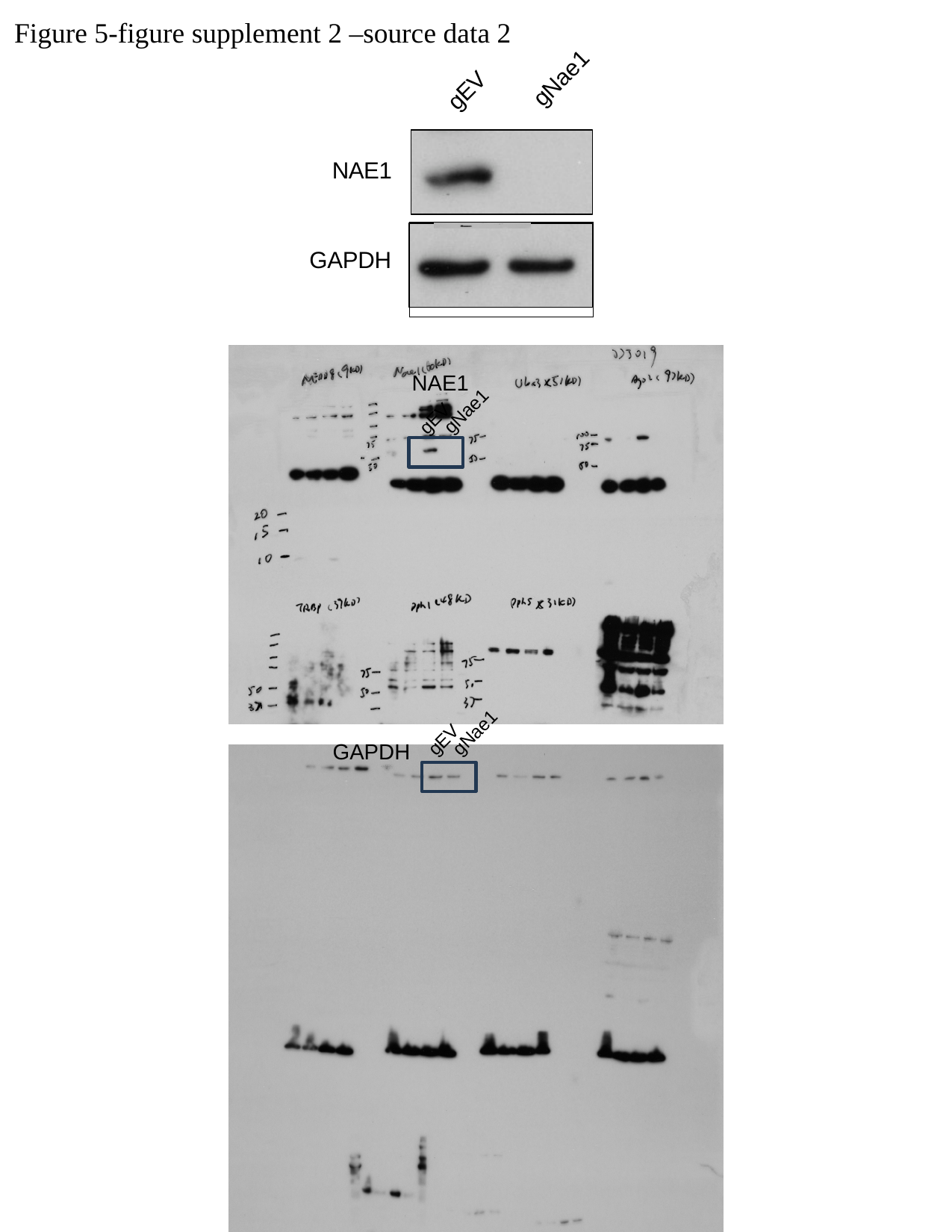

Figure 5-figure supplement 2 –source data 2
gNae1
gEV
NAE1
GAPDH
NAE1
gNae1
gEV
gNae1
gEV
GAPDH

### Slide 7
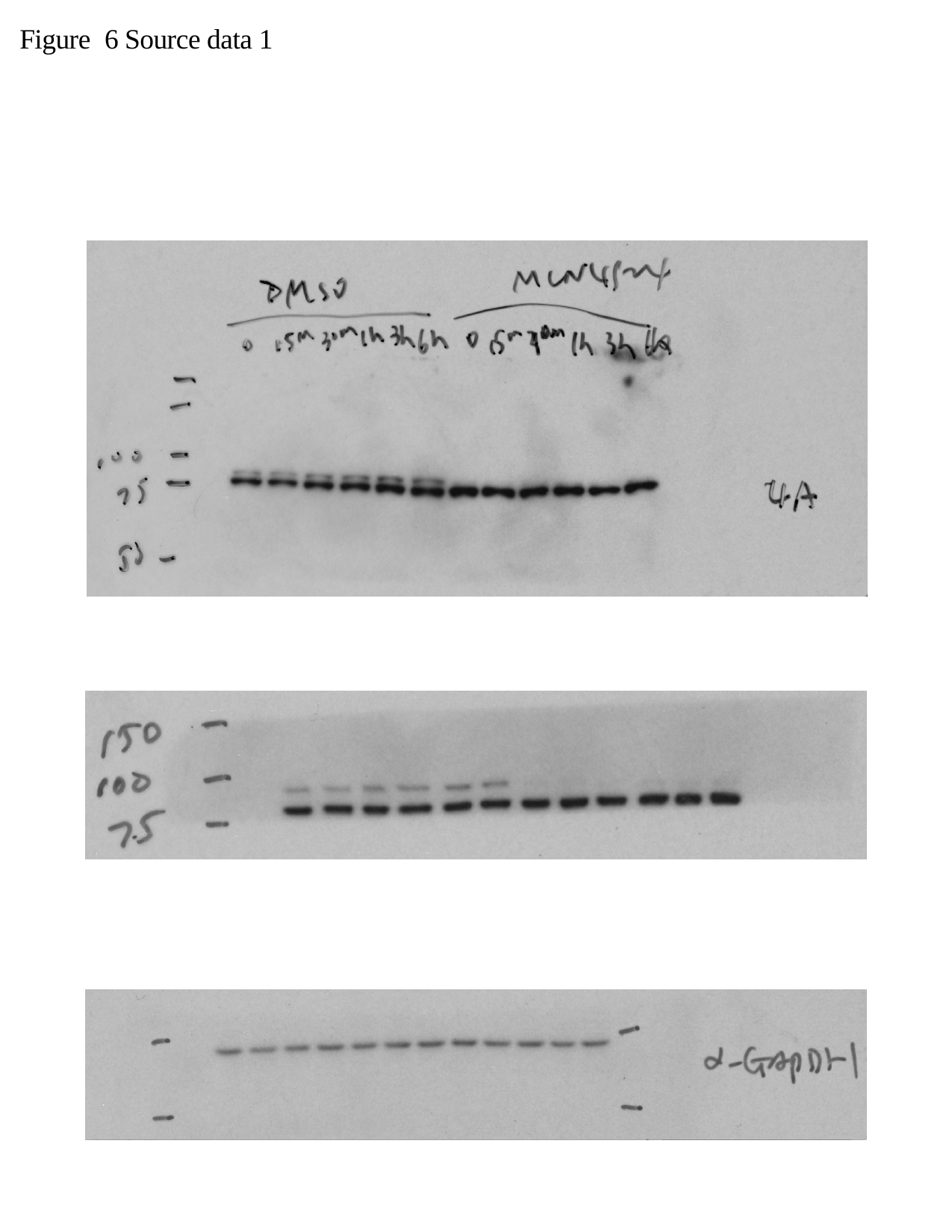

Figure 6 Source data 1

### Slide 8
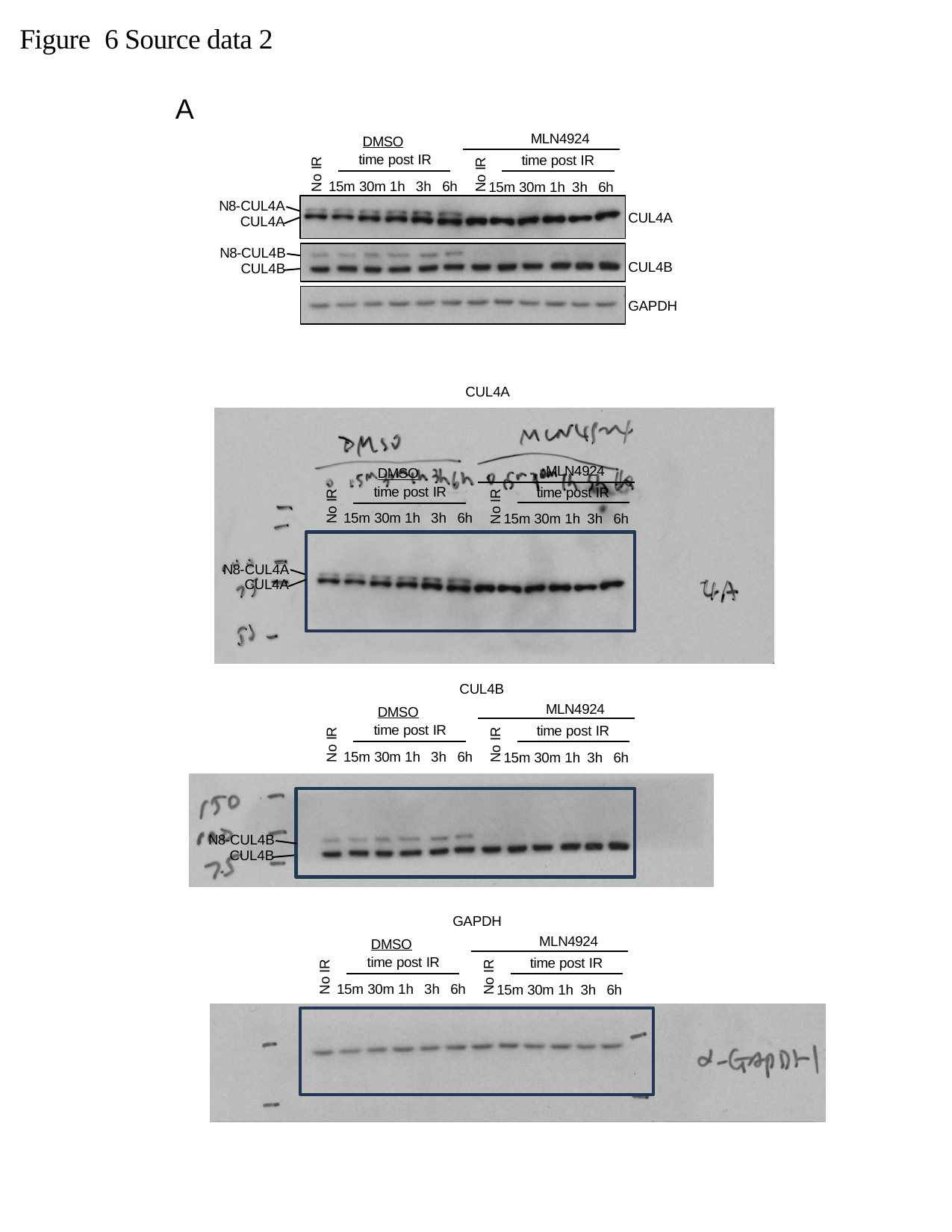

Figure 6 Source data 2
A
MLN4924
	DMSO
time post IR
15m 30m 1h 3h 6h
time post IR
No IR
No IR
15m 30m 1h 3h 6h
N8-CUL4A
CUL4A
CUL4A
N8-CUL4B
CUL4B
CUL4B
GAPDH
CUL4A
MLN4924
	DMSO
time post IR
15m 30m 1h 3h 6h
time post IR
No IR
No IR
15m 30m 1h 3h 6h
N8-CUL4A
CUL4A
CUL4B
MLN4924
	DMSO
time post IR
15m 30m 1h 3h 6h
time post IR
No IR
No IR
15m 30m 1h 3h 6h
N8-CUL4B
CUL4B
GAPDH
MLN4924
	DMSO
time post IR
15m 30m 1h 3h 6h
time post IR
No IR
No IR
15m 30m 1h 3h 6h

### Slide 9
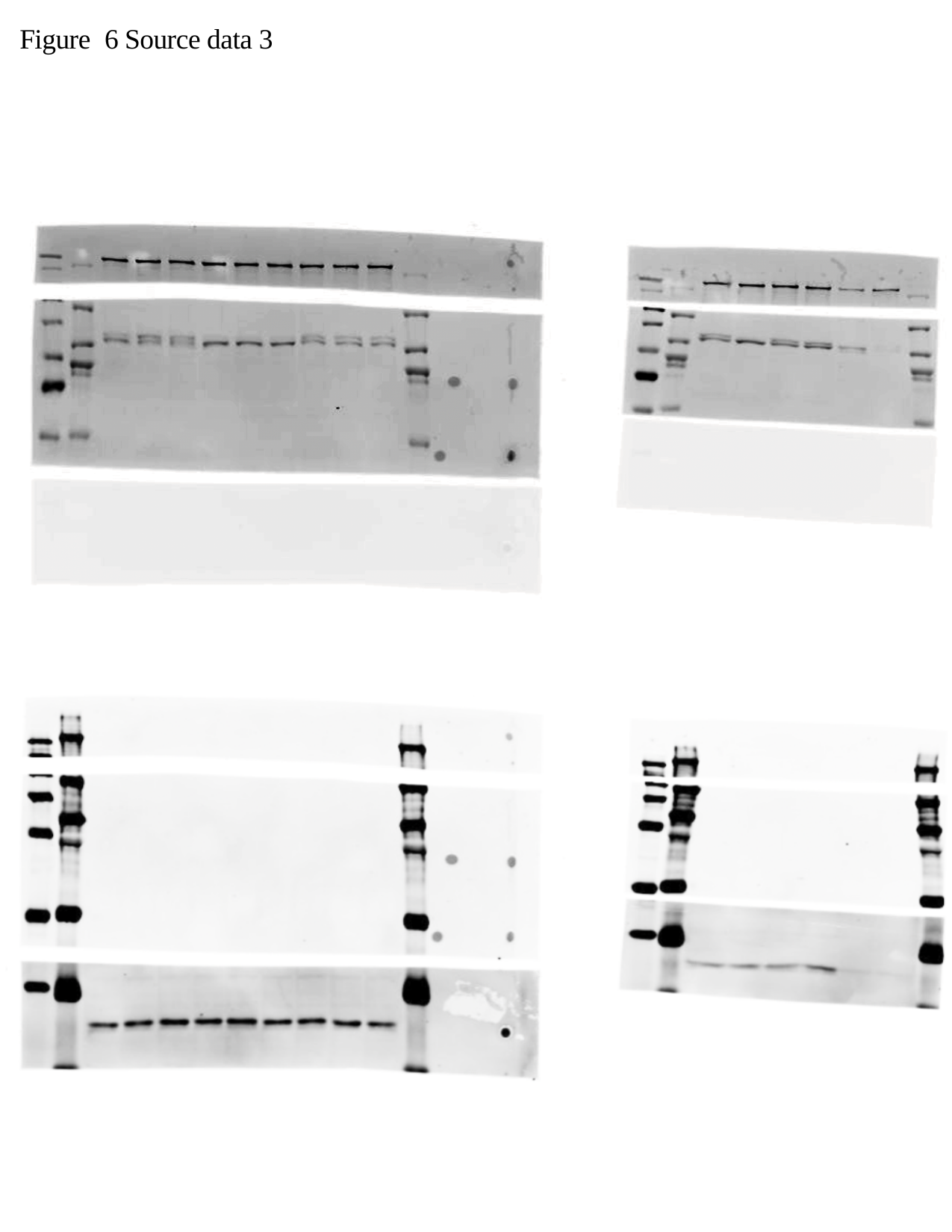

Figure 6 Source data 3

### Slide 10
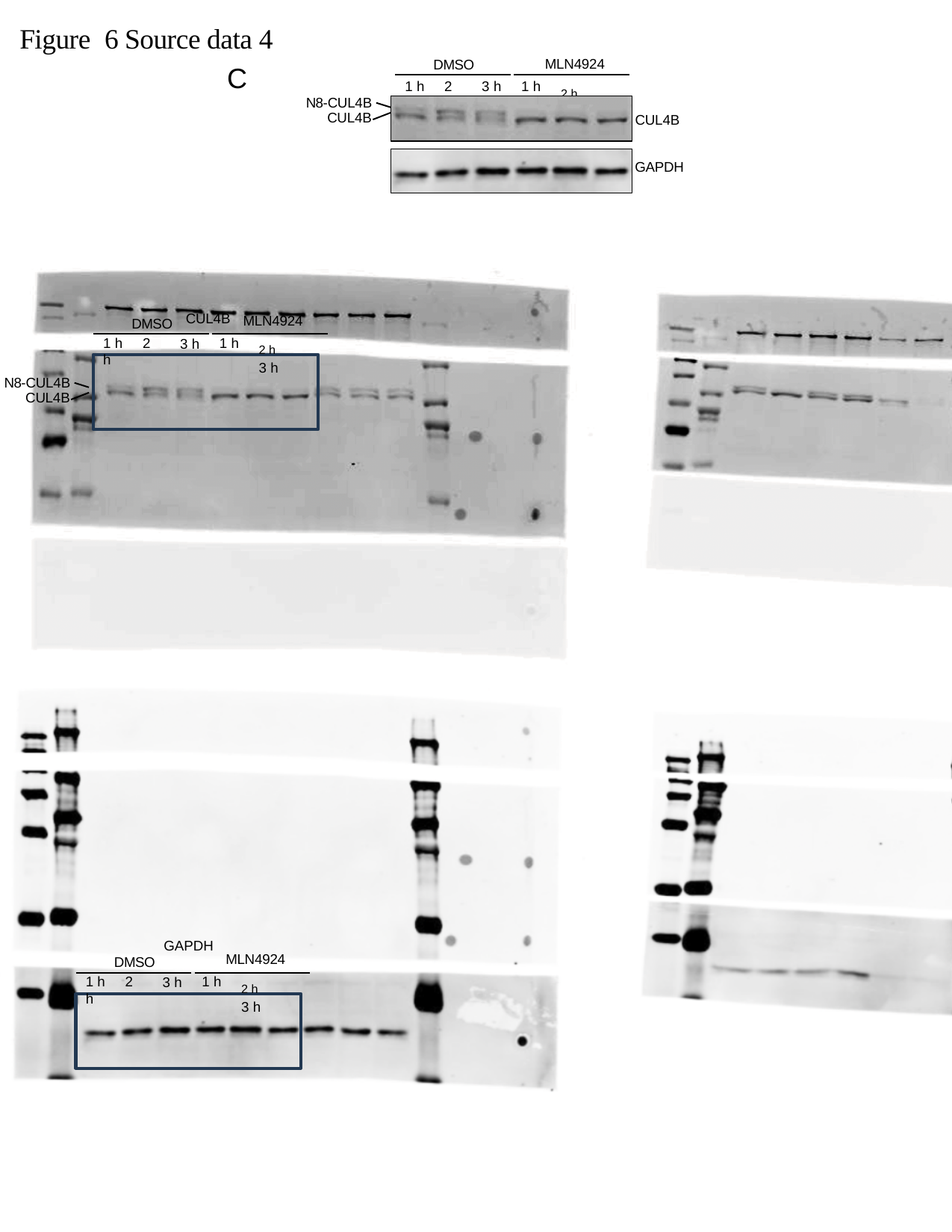

Figure 6 Source data 4
MLN4924
^2^ ^h^	3 h
C
DMSO
1 h	2 h
1 h
3 h
N8-CUL4B
CUL4B
CUL4B
GAPDH
MLN4924
^2^ ^h^	3 h
CUL4B
DMSO
1 h	2 h
1 h
3 h
N8-CUL4B
CUL4B
GAPDH
MLN4924
^2^ ^h^	3 h
DMSO
1 h	2 h
1 h
3 h

### Slide 11
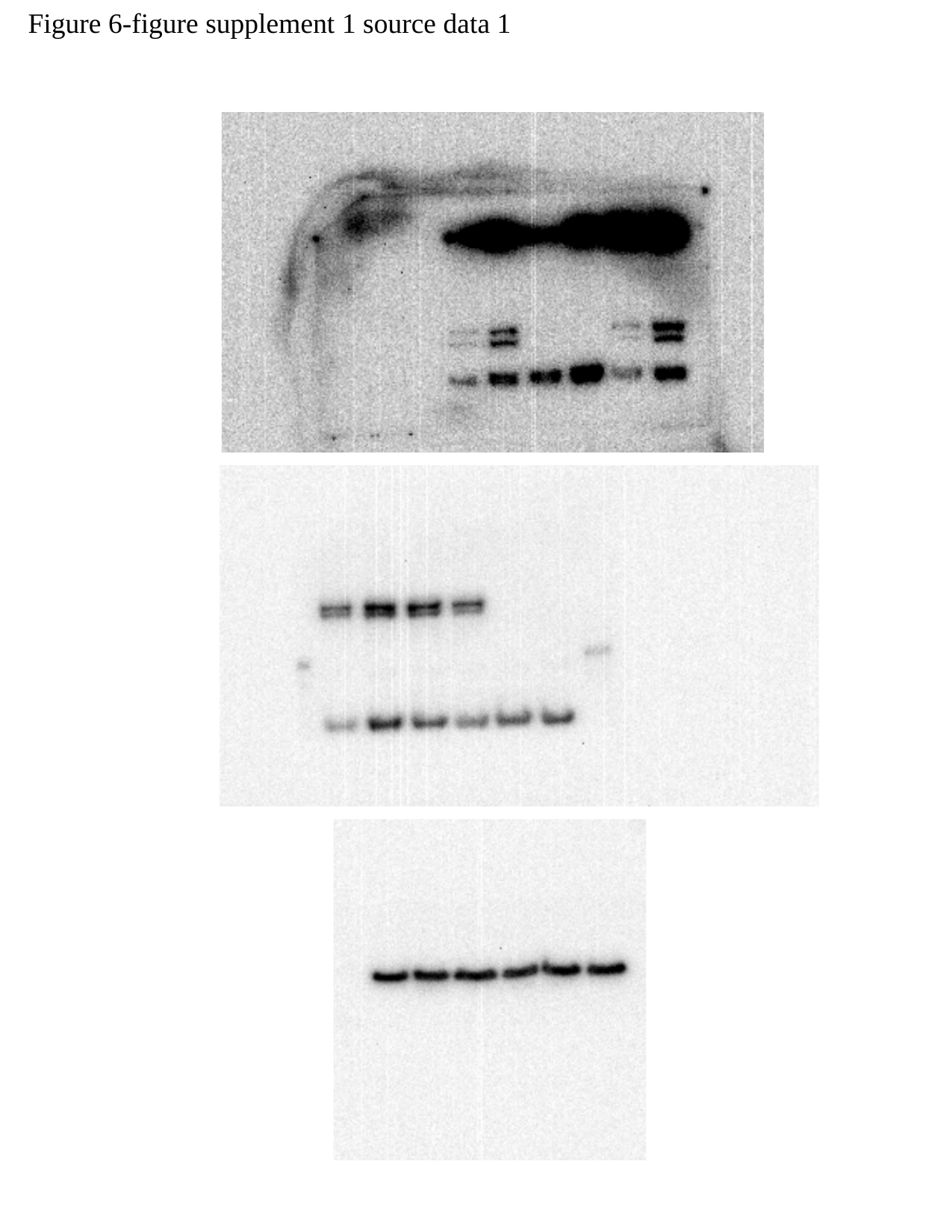

Figure 6-figure supplement 1 source data 1

### Slide 12
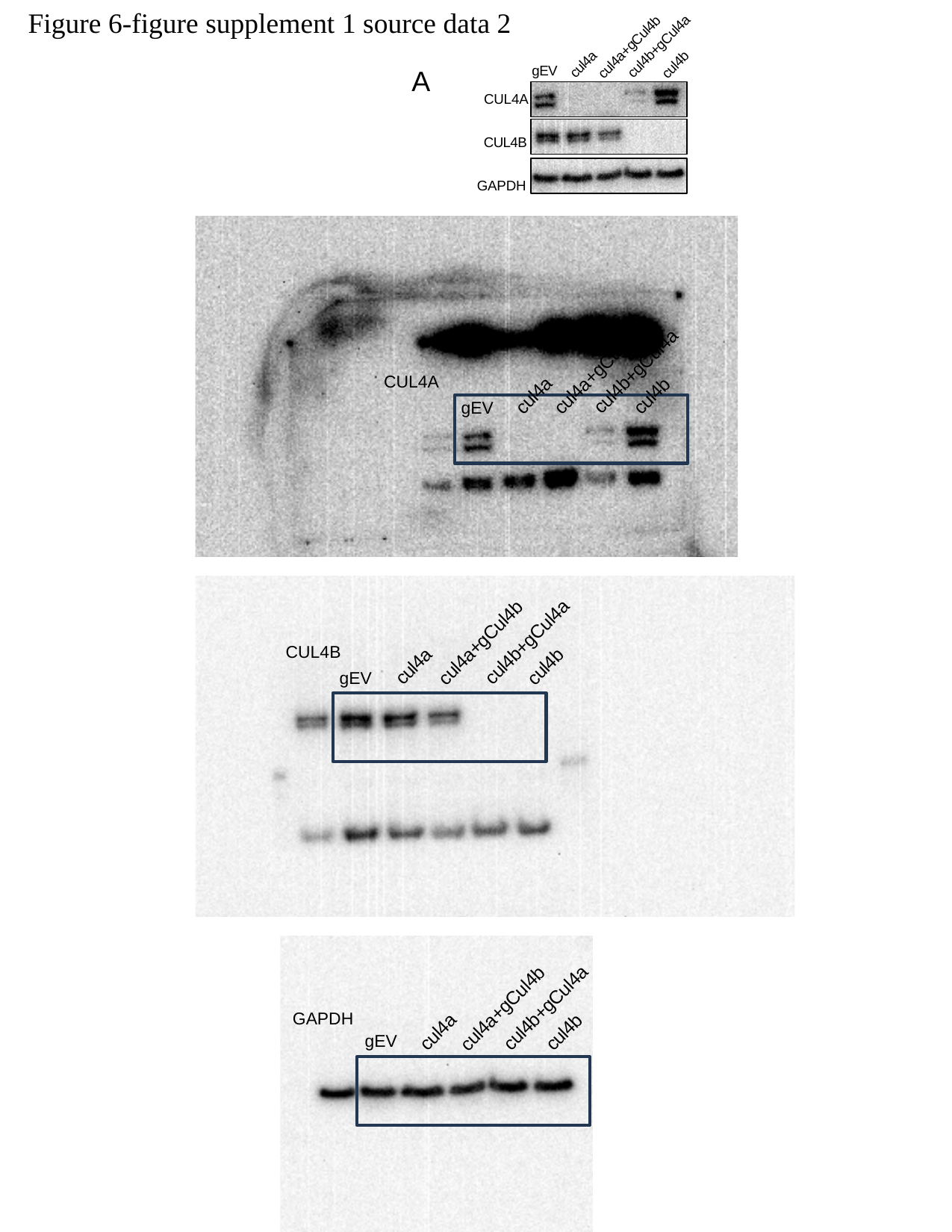

Figure 6-figure supplement 1 source data 2
cul4b+gCul4a
cul4a+gCul4b
cul4a
cul4b
A
gEV
CUL4A
CUL4B GAPDH
cul4a+gCul4b
cul4b+gCul4a
CUL4A
cul4a
cul4b
gEV
cul4a+gCul4b
cul4b+gCul4a
CUL4B
cul4a
cul4b
gEV
cul4a+gCul4b
cul4b+gCul4a
GAPDH
cul4a
cul4b
gEV
